# Pattern-triggered immunity induced by dsRNA against plant viruses involves two distinct branches, one of which depends on *Coronatine insensitive1* (*COI1*)

**DOI:** 10.64898/2026.10.29.762747

**Authors:** Khouloud Necira, Ifigeneia Triantafyllou, Celia González, Tomás Canto, Francisco Tenllado

## Abstract

Double-stranded RNA (dsRNA) is a conserved molecular signature of RNA virus generated during infection that can activate antiviral pattern-triggered immunity (PTI) in plants. However, the effectiveness of dsRNA-induced PTI against viruses from diverse taxa, its impact on different stages of the infection, as well as effectors used by viruses other than tobamoviruses to counteract this defense response remain poorly explored. Here, we established an *Agrobacterium*-mediated transient expression assay for inducing PTI through intracellular production of nonspecific dsRNA in *Nicotiana benthamiana*. Expression of nonspecific dsRNA restricted cell-to-cell movement and accumulation of three RNA viruses representing different taxonomic groups: *Tobacco mosaic virus*, *Turnip mosaic virus*, and *Tobacco rattle virus*. Co-expression of dsRNA together with a P25 movement protein (MP) mutant of *Potato virus X* (PVX) defective in RNA silencing suppression, prevented dsRNA-induced callose accumulation at plasmodesmata (PD) and restored efficient viral cell-to-cell movement. Interestingly, dsRNA also reduced viral RNA accumulation at the cellular level, revealing an impact of dsRNA-based PTI on virus replication independently of cell-to-cell movement. We further showed that the dsRNA-induced antiviral response required jasmonic acid (JA) signaling. Silencing of the JA receptor *Coronatine insensitive1* (*COI1*) abolished dsRNA-induced PD callose deposition and the associated restriction of virus movement and accumulation. Accordingly, dsRNA-induced expression of defense-related genes was compromised in *COI1*-silenced plants. Next, we conducted RNA-seq analysis to identify early biological processes involved in COI1*-*dependent, dsRNA-induced immunity. The differential expression of 198 genes, some potentially involved in the regulation of callose accumulation, in *COI1*-silenced plants suggested that JA signaling contributes to shaping the transcriptional response to nonspecific dsRNA. Together, our findings establish nonspecific dsRNA-induced PTI as a multilayered antiviral response that restricts pathogen replication and cell-to-cell movement, and identify a heterologous MP, PVX P25, as an effector that counteracts the PD component of this defense response in a COI1*-*dependent manner.

**Author summary:** Plants can recognize dsRNA, a molecule produced during infection by RNA viruses, and activate nonspecific immune responses against invading viruses. However, we still know relatively little about how effective this immune response is against distinct viruses and how viruses overcome it. In this study, we develop a method to activate plant immunity against three viruses from different taxa by producing nonspecific dsRNA inside plant cells. We show that dsRNA interfers with virus infection at two different levels: restriction of virus spread between neighboring cells and reduction of virus replication in isolated cells. As a counter-defense strategy, the movement protein of *Potato virus X* hampers dsRNA-induced immunity by preventing the accumulation of callose, a polysaccharide polymer that prevents virus spread between cells. We further show that Coronatine insensitive1, the jasmonic acid receptor involved in regulating plant responses to stress, is required for dsRNA-induced restriction of virus spread. Finally, we identify changes in gene expression associated with this COI1-dependent response to dsRNA. Our findings reveal that plants can defend themselves against invading viruses at different stages of infection, opening new avenues for research into broad-spectrum antiviral strategies for crop protection.

## Introduction

Plant viruses are recognized by their hosts through multiple layers of innate immunity that restrict viral infection at different stages of the infection cycle (Lozano-Durán, 2024; Ngou et al., 2022). In addition to sequence-specific antiviral defenses, plants activate pattern-triggered immunity (PTI) upon perception of conserved molecular signatures associated with invading pathogens, known as pathogen-associated molecular patterns (PAMPs), by pattern recognition receptors (PRRs). Double-stranded RNA (dsRNA), generated either as replication intermediates of RNA viruses or from hairpin RNA structures, has emerged as a PAMP capable of activating antiviral defenses (Chi et al., 2026; Necira et al., 2024; Niehl et al., 2016). Notably, dsRNA-induced antiviral responses can be triggered by both sequence-specific and nonspecific dsRNA, indicating that at least part of the plant response to dsRNA is independent of sequence specificity and therefore distinct from canonical RNA silencing. However, nonspecific dsRNA was considered a less effective inducer of antiviral immunity than sequence-specific dsRNA capable of triggering RNA silencing (Necira et al., 2024; Weiss et al., 2025). Exogenously applied nonspecific dsRNA or its synthetic analog polyinosinic-polycytidilic acid (poly (I:C)) has been shown to induce defense responses and restrict infection by diverse plant viruses, although the mechanisms underlying its perception and downstream antiviral effects remain incompletely understood (Heinlein, 2025).

Antiviral PTI responses induced by dsRNA share several features with antimicrobial PTI, including the induction of mitochondrial reactive oxygen species (ROS), ethylene production, activation of mitogen-activated protein kinases, and expression of defense-related genes (Chi et al., 2026; Nicaise and Candresse, 2017; Niehl et al., 2016; Yuan *et al*., 2021). However, poly(I:C) perception did not trigger apoplastic ROS production, suggesting that dsRNA-induced PTI engages signaling pathways that differ from those activated by microbial PAMPs (Huang et al., 2023). In *Arabidopsis thaliana*, a signaling pathway involving the PRR coreceptor kinase SERK1, the receptor-like cytoplasmic kinases BIK1/PBL1, the plasmodesmal proteins PDLP1, PDLP2, and PDLP3, calmodulin-like protein 41 (CML41), and calcium signaling has been implicated in the dsRNA-based PTI response (Huang et al., 2023). More recently, it was reported that the leucine-rich repeat protein AtCERES and the mitochondrial protein AtXIAN mediate dsRNA-triggered plant immune responses, although a direct binding of AtCERES to dsRNA was not demonstrated (Chi et al., 2026).

One important feature of dsRNA-induced PTI is the reinforcement of intercellular barriers that restrict virus movement through plasmodesmata (PD) (Huang et al., 2023). PD are dynamic intercellular channels that regulate the movement of macromolecules between adjacent cells and represent a major route exploited by viruses for cell-to-cell spread (Burch-Smith, 2026). Plants regulate PD permeability in response to developmental and environmental cues as well as pathogen invasion, with deposition of β-1,3-glucan (callose) at the neck regions of PD. Many genes related to callose synthesis (β-1,3-glucan synthases) or degradation (β-1,3-glucanases) as well as PD-localized callose binding proteins (PDCBs) among others, have been identified as crucial regulators of symplasmic connectivity. Increased callose accumulation reduces the PD size exclusion limit and thereby restricts the intercellular trafficking of viral genomes and movement complexes. Accordingly, the regulation of PD permeability represents an important component of plant antiviral immunity and a major target of viral counter-defense strategies (Burch-Smith, 2026). Recent studies have provided evidence that movement proteins (MPs) of tobamoviruses (genus *Tobamovirus*, family *Virgaviridae*) suppress dsRNA-induced callose deposition at PD, allowing viruses to overcome this PD-associated antiviral defense (Huang et al., 2023). These findings and others indicate that viral MPs may function not only as components of the viral movement machinery but also as effectors that actively interfere with host immune responses (Huang et al., 2023; Kong *et al*., 2018). However, evidence that viral MPs suppress dsRNA-induced PD closure remains largely restricted to tobamoviruses, and it is unclear whether this represents a mechanism broadly conserved among plant viruses with distinct cell-to cell movement strategies. Furthermore, the effects of dsRNA-induced PTI on other steps of the viral infection cycle, e.g., virus replication, have been poorly explored (Huang et al., 2023; Necira *et al*., 2024). In bacterial and fungal pathosystems, plant immunity can restrict pathogens at multiple stages of infection, including host entry, establishment, proliferation, and dissemination (Jian et al., 2024).

*Potato virus X* (PVX), a member of the genus *Potexvirus* in the family *Alphaflexiviridae*, encodes a triple gene block (TGB) module in which TGB1 (P25) plays a central role in the cell-to-cell movement of viral RNA (Verchot, 2022). P25 is an RNA-binding protein with helicase activity that associates with viral RNA and increases the size exclusion limit of PD, thereby facilitating the passage of viral movement complexes between adjacent cells (Howard et al., 2004; Park et al., 2014). TGB2 and TGB3 cooperate with P25 to target the viral movement machinery to PD. In addition to its role in virus movement, P25 functions as a suppressor of antiviral RNA silencing (Voinnet et al., 2000). Thus, distinguishing its effects on RNA silencing from those associated with PD regulation is essential when investigating a potential role of PVX P25 in the suppression of dsRNA-induced PTI.

Phytohormones are key regulators of plant immunity, functioning as signaling molecules that integrate and coordinate immune responses to diverse environmental and biotic stresses. In particular, antimicrobial PTI is regulated by phytohormones, with salicylic acid (SA) and jasmonic acid (JA) acting as key regulators of the immune signaling network (Zhang et al., 2025). For example, application of bacterial RNA has been shown to activate PTI against *Pseudomonas syringae* in Arabidopsis, a response that requires both SA- and JA-dependent signaling (Lee et al. 2016). In addition, extracellular self-DNA has been reported to inhibit growth and activate defense responses in Arabidopsis and tomato, supporting the hypothesis that it functions as a damage-triggered immune response that depends on JA signaling (Zhou et al., 2023). However, the contribution of phytohormone signaling to dsRNA-mediated PTI during viral infection remains unexplored.

Using an *Agrobacterium*-mediated transient expression assay to deliver nonspecific dsRNA in *Nicotiana benthamiana*, we examined the effects of PVX P25 on dsRNA-induced callose deposition, virus accumulation, and cell-to-cell movement of three RNA viruses representing distinct taxonomic groups. In addition, we investigated the ability of nonspecific dsRNA to interfere with the replication of these viral amplicons in agropatch assays and protoplasts. Our results support the existence of two distinct branches in antiviral PTI induced by dsRNA. PVX P25 suppressed dsRNA-induced callose accumulation and counteracted the associated restriction of virus cell-to-cell movement, but it did not overcome dsRNA-mediated inhibition of viral replication. We further investigated the contribution of JA and SA signaling to dsRNA-induced antiviral PTI. We found that COI1-dependent JA signaling is involved in the regulation of dsRNA-induced callose deposition and the restriction of virus cell-to-cell movement. Taken together, our findings suggest that viral movement proteins act as broad-range suppressors of dsRNA-induced PTI and provide evidence that nonspecific dsRNA activates antiviral defenses that operate at both cellular and tissue levels.

## Results

### PTI-based antiviral defense induced by *Agrobacterium*-mediated transient expression of dsRNA is suppressed by PVX MP

The MP of *Tobacco mosaic virus* (TMV) suppress dsRNA-induced PTI by disrupting callose deposition at PD, thereby facilitating virus movement (Huang et al., 2023). However, current evidence for the ability of MPs to inhibit dsRNA-induced PD closure is limited to tobamovirus MPs. To assess whether another viral MP is capable of suppressing dsRNA-induced PTI in a heterologous system, we used two single-amino-acid mutants of the PVX MP, P25 T117A and K124E, with distinctive properties. These mutants are defective in RNA silencing suppression, thereby avoiding any influence of the suppression of host’s antiviral RNA silencing on the interpretation of the PTI assays (Bayne et al., 2005). We confirmed previous results showing that mutants T117A and K124E were deficient in suppressing the silencing of the GFP reporter in leaf patch assays, whereas P25 wt was active in this function (S1 Fig).

Leaf patches of *N. benthamiana* plants were infiltrated with *Agrobacterium* cultures containing binary vectors expressing either (-)ssRNA or hairpin RNA (hereafter dsRNA) sequences derived from *Pepper mild mottle virus* (PMMoV) together with agrobacteria expressing mutant versions of P25 or GUS as a control. After 3 days, leaf patches were inoculated with leaf sap derived from plants infected with TMV-GFP, and virus infection was monitored by detecting GFP foci. At 4 days post-inoculation (dpi), there was a significant reduction (2.2-fold) in the size of GFP foci in patches expressing dsRNA + GUS compared to (-)ssRNA + GUS-expressing patches. By contrast, treatments with either dsRNA + P25 T117A or dsRNA + P25 K124E did restore the size of GFP foci to that in (-)ssRNA + GUS-expressing patches (Fig. 1A).

**Fig 1.**
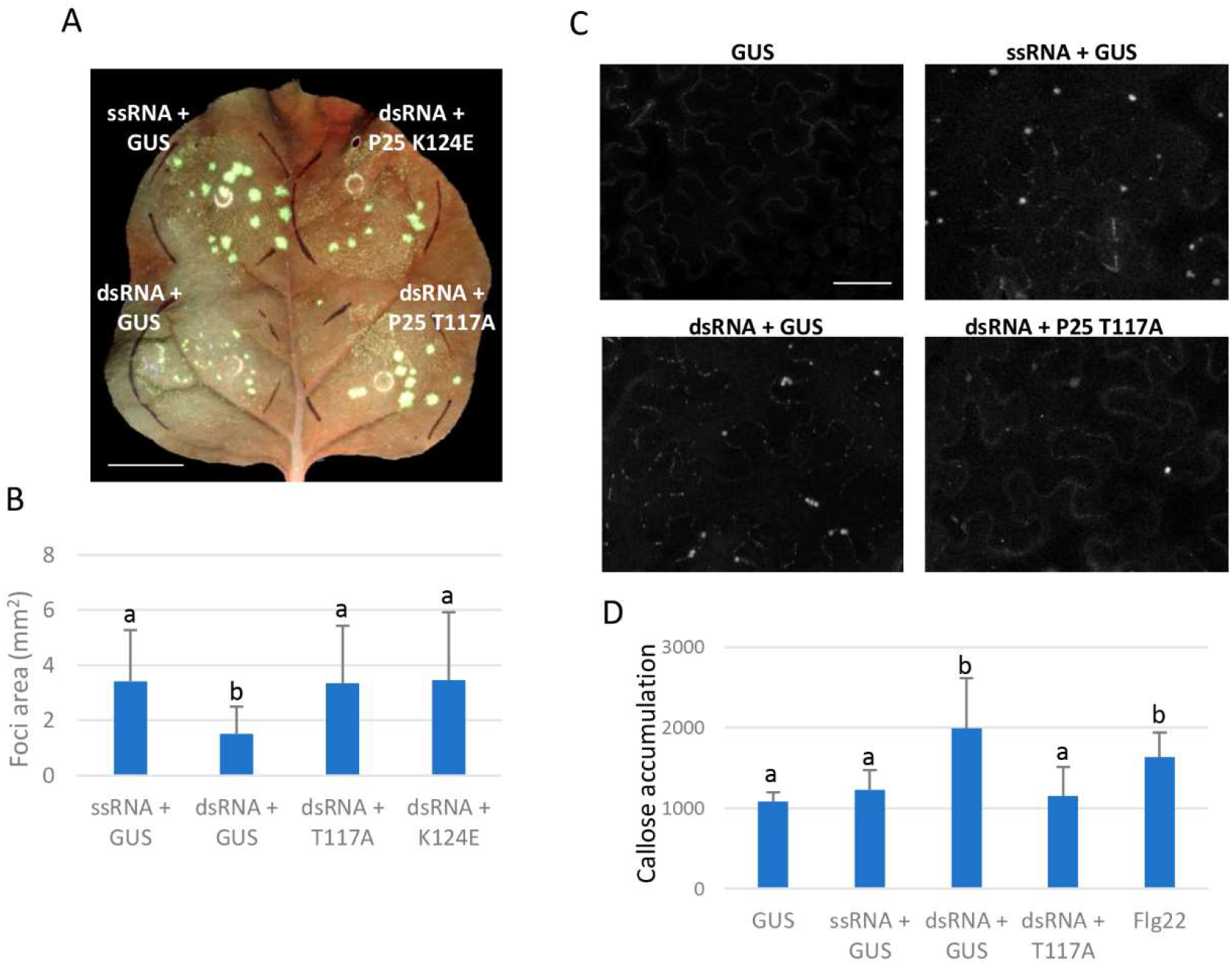
PVX P25 alleviates dsRNA-induced restriction of TMV-GFP cell-to-cell movement by preventing callose accumulation. (A) Leaves of *N. benthamiana* were infiltrated with combinations of *Agrobacterium* cultures containing binary constructs expressing dsRNA sequences derived from PMMoV plus either GUS, T7-tagged versions of P25 T117A or K124E, and expressing (-)ssRNA plus GUS, as indicated. After 3 days, the infiltrated patches were inoculated with TMV-GFP. The inoculated leaves were examined under UV light at 4 days post-inoculation (dpi). Scale bar denotes 1 cm. (B) Mean numbers ± standard deviations of infection foci on infiltrated patches with the different treatments at 4 dpi. Twelve patches from 6 plants for each treatment were analyzed. Different letters indicate significant differences determined by employing Scheffé’s test (P < 0.05). (C) Callose spots at plasmodesmata (PD) were visualized upon aniline blue staining of epidermal cells on patches expressing GUS alone or combinations of dsRNA plus either GUS or P25 T117A and of (-)ssRNA plus GUS, in the absence of viral infection. The bacterial PTI elicitor derived from flagellin (flg22) was used as a control. Scale bar, 20 μm. (D) Relative PD callose content in the different treatments mentioned above. The mean values of callose intensities in individual PD (>200) measured in three leaf discs taken from three independent biological replicates per each treatment. Different letters indicate significant differences determined by employing Dunnett’s T3 test (P < 0.05). Experiments of callose deposition were repeated once more with similar results.

To determine whether *Agrobacterium*-mediated transient expression of dsRNA triggers callose accumulation in infiltrated patches, in vivo aniline blue staining was performed on patches expressing GUS, (-)ssRNA, dsRNA, the combination dsRNA + P25 T117A or treated with the bacterial PTI elicitor flg22. In agreement with the reduction in the size of TMV-GFP foci observed above (Fig. 1A), callose accumulation at PD was increased by 63% in patches treated with dsRNA compared to (-)ssRNA-expressing patches (Fig. 1C). Similarly, increased callose deposition at PD was also observed in patches treated with flg22. Importantly, the P25 T117A mutant protein derived from PVX reduced the levels of callose deposition induced by dsRNA to control levels. These results indicate that the inhibition of TMV-GFP cell-to-cell movement induced by transient expression of nonspecific dsRNA was associated with enhanced callose deposition at PD, which was prevented by expression of a heterologous MP, the PVX P25.

In a subsequent experiment, the effect of the different treatments on the accumulation level of TMV-GFP was measured in the inoculated patches at 5 dpi, a time point that permits a more reliable quantification of viral load (Fig. 2A). The combination (-)ssRNA + P25 T117A was analyzed as an additional control. Virus accumulation was lower in leaf patches treated with dsRNA + GUS than in (-)ssRNA + GUS or (-)ssRNA + P25 T117A-expressing patches as assayed by western blot analysis, whereas TMV-GFP accumulation was partially restored in treatment with dsRNA + P25 T117A (Fig. 2B). We confirmed that P25 T117A mutant protein accumulated in patches infiltrated with dsRNA + P25 T117A at levels comparable to those in (-)ssRNA + P25 T117A-expressing patches (Fig. 2C). Comparative analysis by quantitative RT-PCR (qRT-PCR) of RNA extracted from agroinfiltrated patches revealed that the level of TMV-GFP RNA in patches treated with dsRNA + GUS was reduced 2.1-fold compared to (-)ssRNA + GUS-expressing patches, and that virus accumulation was restored to control levels in patches expressing dsRNA + P25 T117A (Fig. 2D). qRT-PCR analysis confirmed transient expression of the (-)ssRNA and dsRNA sequences, using primers that amplify the spacer sequence (Phe) between the arms of the T-DNA constructs (Fig. 2E). Note that dsRNA expression reached similar levels in both dsRNA + GUS and dsRNA + P25 T117A-expressing patches, whereas (-)ssRNA accumulated at lower levels probably because it was more susceptible to degradation by cellular nucleases.

**Fig 2.**
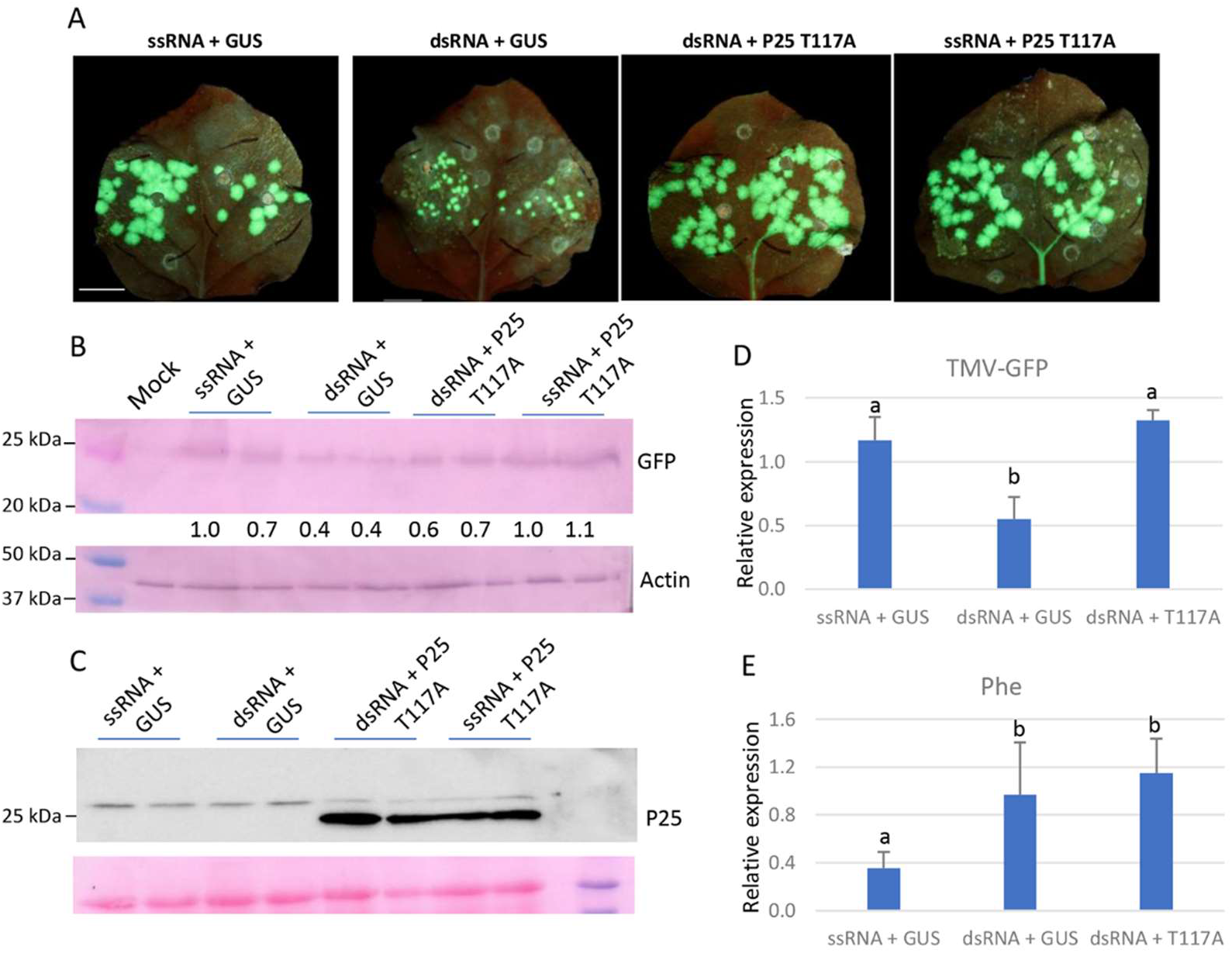
PVX P25 interferes with dsRNA-induced PTI. (A) Leaves of *N. benthamiana* were infiltrated with combinations of *Agrobacterium* cultures containing binary constructs expressing dsRNA sequences derived from PMMoV plus either GUS or T7-tagged P25 T117A, and expressing (-)ssRNA plus either GUS or T7-tagged P25 T117A, as indicated. After 3 days, the infiltrated patches were inoculated with TMV-GFP. The inoculated leaves were examined under UV light at 5 days post-inoculation (dpi). Scale bar denotes 1 cm. (B) Western blot analysis of extracts derived from leaf patches at 5 dpi, using antibodies against GFP (upper panel) or actin (lower panel), as a loading control. The intensity of the GFP bands, normalized for the loading controls, was quantified by densitometry analysis. The value for the band in leaf patches infiltrated with (-)ssRNA plus GUS was set at 1 and other data calculated relative to this value. (C) Western blot analysis of extracts derived from the different treatments mentioned above, using antibodies against the T7 epitope. The panel below the western blot shows the Ponceau S-stained membrane after blotting. (D,E) qRT-PCR was used to analyze the accumulation of TMV-GFP RNA (D) and the phleomycin (Phe) RNA sequence (E) used as a spacer between the arms of the T-DNA constructs at 5 dpi. Expression of the 18S rRNA gene served as a control. Data represent the means ± standard deviation of at least 3 replicates, each consisting of a pool of 6 plants that received the same treatment. Different letters indicate significant differences determined by employing Mann-Whitney U test (D) and Duncan’s test (E) (P < 0.05).

To determine if the PVX MP also interferes with dsRNA-induced PTI against other viruses, we tested the effect of dsRNA-transient expression on infection by the potyvirus *Turnip mosaic virus* (TuMV) and the tobravirus *Tobacco rattle virus* (TRV), either alone or co-expressed with P25 T117A. There was a significant reduction in the size of TuMV-GFP and TRV-GFP foci in patches expressing dsRNA + GUS compared to (-)ssRNA + GUS-expressing patches, which was restored in the treatment with dsRNA + P25 T117A (S2 Fig). Consistently, TRV-GFP RNA levels were 2.3-fold lower in dsRNA-expressing patches than in (-)ssRNA-expressing patches at 2 dpi, whereas co-expression of dsRNA with P25 T117A restored viral RNA levels to those observed in (-)ssRNA + GUS-expressing patches, as determined by qRT-PCR.

### dsRNA-mediated PTI restricts viral replication

To determine whether dsRNA-induced PTI affects other steps of virus infection besides cell-to-cell movement, we co-agroinfiltrated a binary vector expressing TuMV-GFP together with binary vectors expressing GUS, (-)ssRNA, dsRNA, and P25 T117A in any of the four combinations mentioned above. The treatment dsRNA + GUS caused a reduction in GFP fluorescence intensity, a proxy for viral replication and accumulation, compared to (-)ssRNA+ GUS and (-)ssRNA + P25 T117A, which was not reverted by P25 T117A expression (dsRNA + P25 T117A) (Fig. 3A). In the absence of a substantial contribution of virus cell-to-cell movement, TuMV-GFP RNA accumulated at lower levels (6.7-fold) in patches expressing dsRNA than in those expressing (-)ssRNA at 4 dpi, as measured by qRT-PCR (Fig. 3B). The combination dsRNA + P25 T117A did not restore the level of accumulation of TuMV-GFP to those observed in the (-)ssRNA + GUS and (-)ssRNA + P25 T117A combinations. Thus, although the PVX MP counteracts dsRNA-induced inhibition of virus cell-to-cell movement, it cannot overcome dsRNA-mediated inhibition of viral replication.

**Fig 3.**
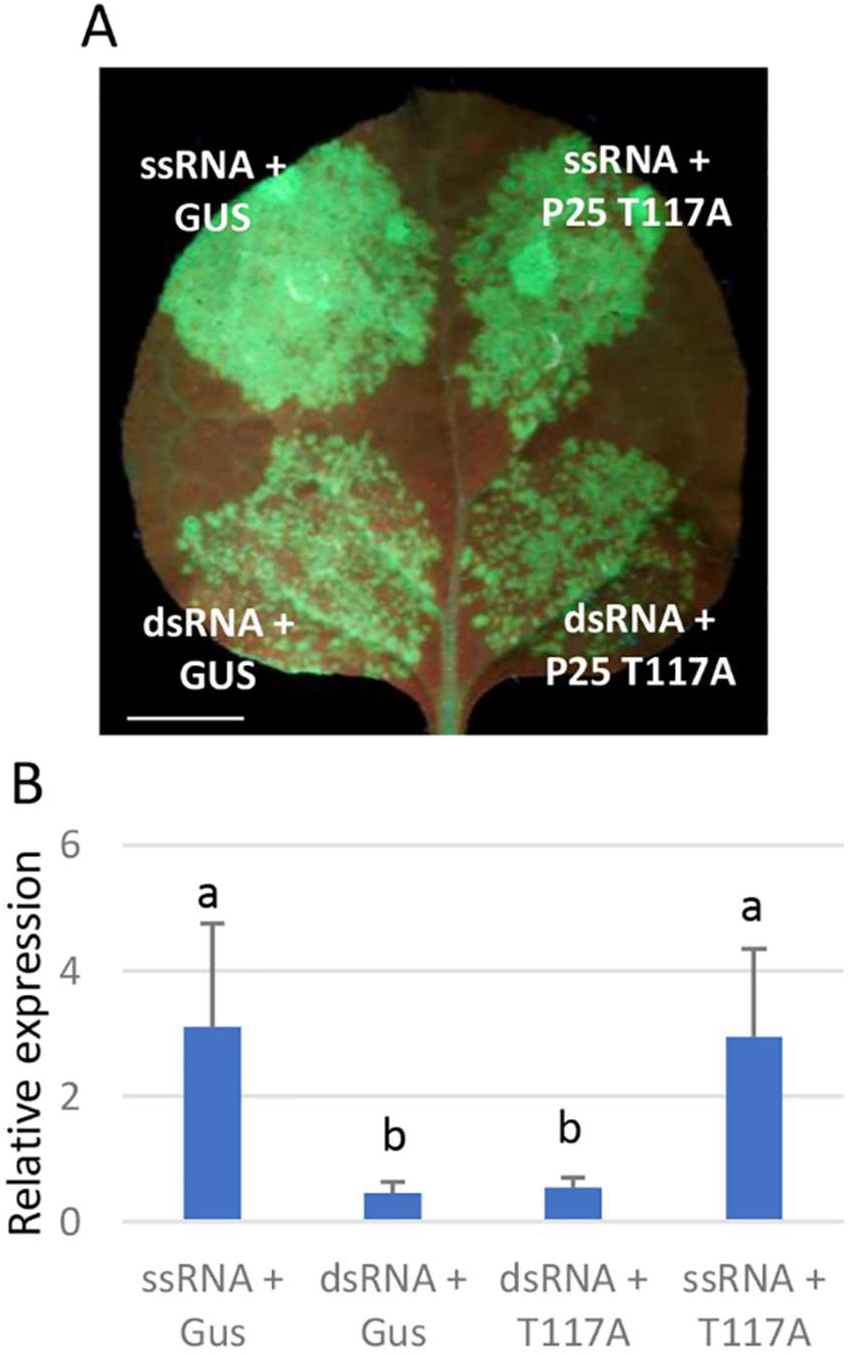
dsRNA-induced PTI restricts TuMV-GFP replication. (A) Leaves of *N. benthamiana* were infiltrated with *Agrobacterium* cultures containing binary constructs expressing TuMV-GFP together with cultures expressing dsRNA sequences derived from PMMoV plus either GUS or T7-tagged P25 T117A, and expressing (-)ssRNA plus either GUS or T7-tagged P25 T117A, as indicated. The inoculated leaves were examined under UV light at 4 days post-inoculation (dpi). Scale bar denotes 1 cm. (B) qRT-PCR was used to analyze the accumulation of TuMV-GFP RNA at 4 dpi. Expression of the 18S rRNA gene served as a control. Data represent the means ± standard deviation of at least 3 replicates, each consisting of a pool of 6 plants that received the same treatment. Different letters indicate significant differences determined by employing Tukey’s test (P < 0.05). The experiment was repeated once more with similar results.

To determine whether the inhibitory effect of dsRNA-induced PTI on viral replication extends to other viruses, we co-agroinfiltrated a binary vector expressing TRV-GFP together with binary vectors expressing one of the following PMMoV-derived sequences: (-)ssRNA, (+)ssRNA, or dsRNA. In addition, we co-agroinfiltrated binary vectors expressing TRV-derived (-)ssRNA and (+)ssRNA, which are complementary and can therefore form dsRNA homologous to the target virus. Two days after virus inoculation, GFP fluorescence intensity and virus accumulation were reduced in patches expressing PMMoV dsRNA compared to those expressing PMMoV (-)ssRNA or (+)ssRNA (Fig. 4). At later observation dates (3 dpi), differences in GFP fluorescence intensity and virus accumulation between PMMoV dsRNA-expressing patches and (-)ssRNA or (+)ssRNA-expressing patches were less pronounced. By contrast, leaf patches expressing both the (-)ssRNA and (+)ssRNA forms of TRV remained virus-free throughout the experiment, indicating that dsRNA homologous to the target virus confers stronger inhibition of TRV-GFP replication than nonspecific dsRNA.

**Fig 4.**
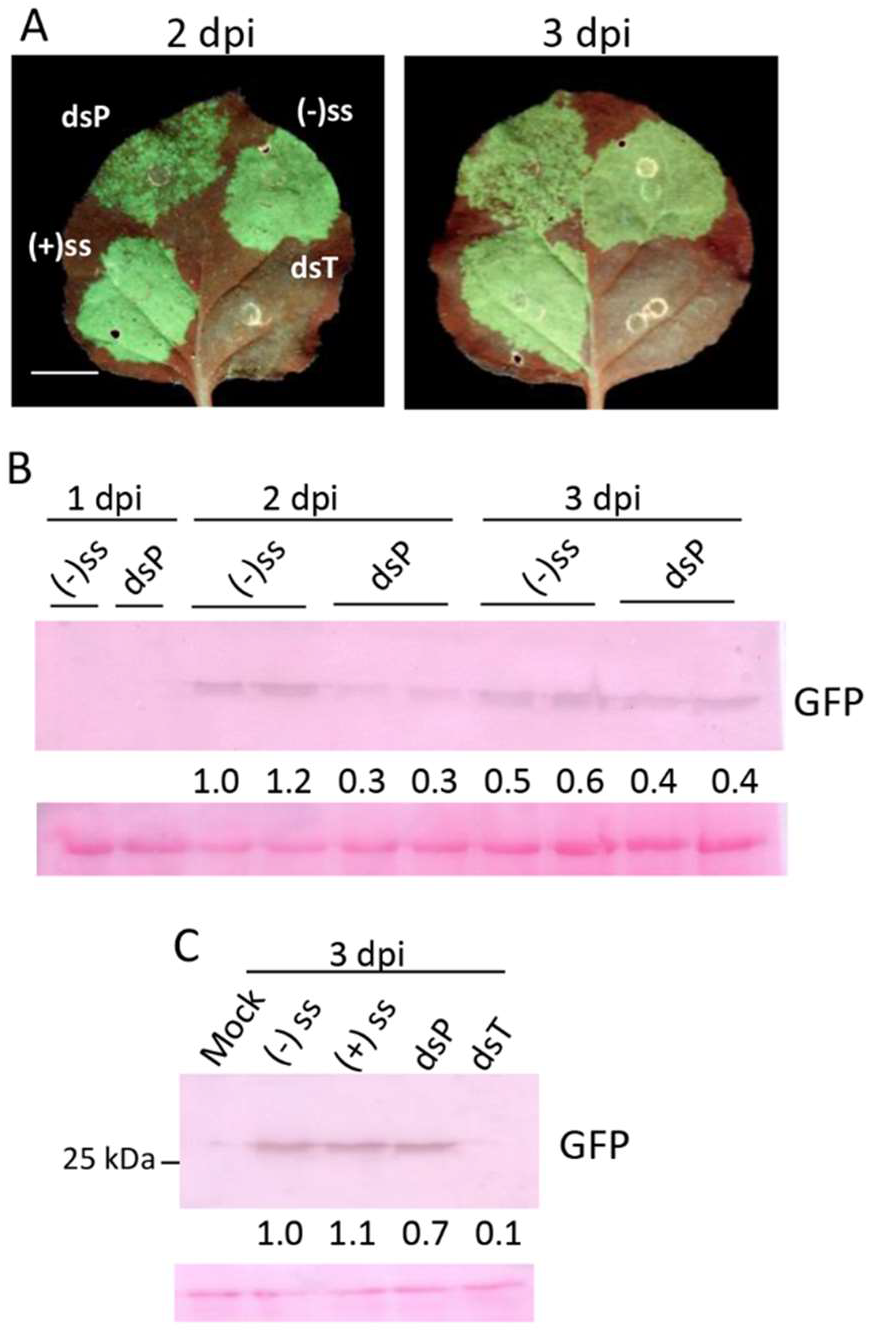
dsRNA-induced PTI restricts TRV-GFP replication. (A) Leaves of *N. benthamiana* were infiltrated with *Agrobacterium* cultures containing binary constructs expressing TRV-GFP together with cultures expressing PMMoV-derived dsRNA (dsP), (-)ssRNA, or (+)ssRNA, and expressing (-)ssRNA plus (+)ssRNA from TRV (dsT), as indicated. The inoculated leaves were examined under UV light at 2 (left panel) and 3 (right panel) days post-inoculation (dpi). Scale bar denotes 1 cm. (B,C) Western blot analyses of extracts derived from leaf patches at 1 to 3 (B) and 3 (C) dpi, using antibodies against GFP. The panels below the western blots show the Ponceau S-stained membrane after blotting. The intensity of the GFP bands, normalized for the loading controls, was quantified by densitometry analysis. The value for the band in leaf patches infiltrated with (-)ssRNA was set at 1 and other data calculated relative to this value.

We then tested whether dsRNA could induce an antiviral PTI response in protoplasts isolated from *N. benthamiana*. We synthesized dsRNA against a 902-bp-long fragment corresponding to the coat protein gene of *Potato virus Y* (dsPVY) and cotransfected it with pTuMV-GFP into protoplasts. We found that cotransfection of pTuMV-GFP with dsPVY significantly reduced TuMV-GFP RNA accumulation (3.2-fold) compared to cotransfection with ssPVY, as determined by qRT-PCR (Fig. 5A). To further substantiate this finding, protoplasts were co-transfected with pTRV RNA1 and either dsPVY or ssPVY. TRV RNA1 levels were reduced by 1.83-fold in cotransfection with dsPVY compared to cotransfection with ssPVY (Fig. 5B). Based on these data, we concluded that nonspecific dsRNA-induced PTI restricts virus replication at the cellular level.

**Fig 5.**
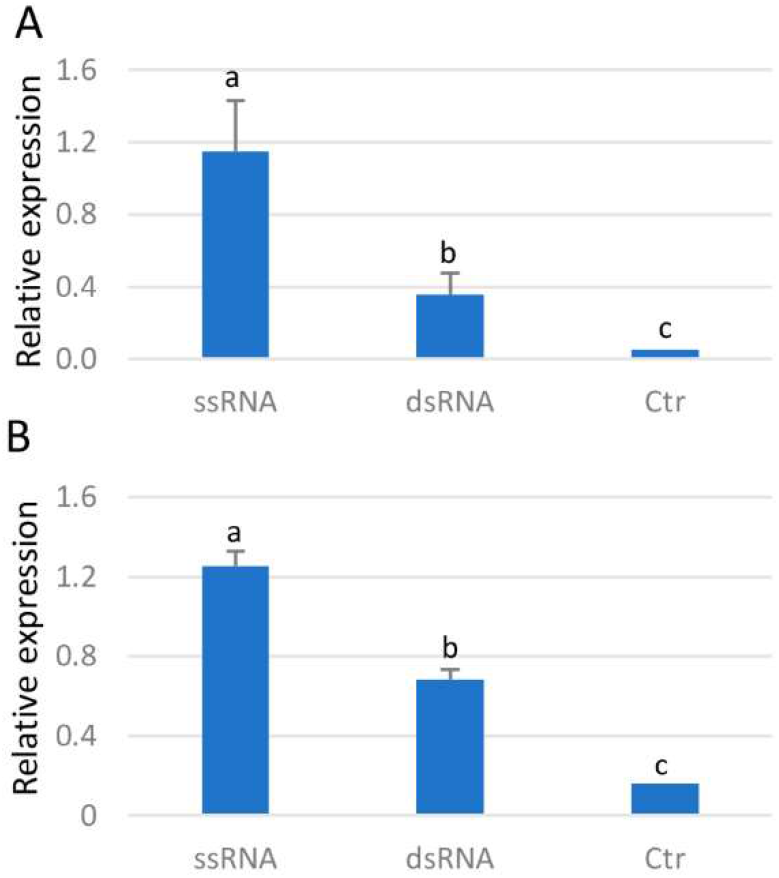
dsRNA-induced PTI restricts virus replication in protoplasts. (A) Protoplasts were co-transfected with plasmid expressing TuMV-GFP together with either in vitro synthetized dsRNA or (-)ssRNA from PVY. qRT-PCR was used to analyze the accumulation of TuMV-GFP RNA at 24 hours post transfection (hpt). (B) Protoplasts were co-transfected with plasmid expressing TRV RNA1 together with either in vitro synthetized dsRNA or (-)ssRNA from PVY. qRT-PCR was used to analyze the accumulation of TRV RNA1 at 24 hpt. Expression of the 18S rRNA gene served as a control. Data represent the means ± standard deviation of at least 3 replicates. Different letters indicate significant differences determined by employing Tukey’s test (*p* < 0.05). Ctr, mock-transfected protoplasts. Experiments were repeated once more with similar results

### PTI-based antiviral defense is dependent on JA signaling

To shed light on the contribution of phytohormone signaling to dsRNA-mediated PTI against infecting viruses, *N. benthamiana* transgenic plants deficient in JA perception (*IRCOI1*) or SA accumulation (*NahG*), as well as wt plants, were used. *IRCOI1* plants stably expressing a hairpin RNA from the JA receptor gene *Coronatine insensitive1* (*COI1*), were partially characterized at both phenotypical and molecular level. *IRCOI1* plants exhibited slightly retarded growth, with occasional rolling of the leaf blades into a cup-shaped form (S3A Fig). qRT-PCR analysis confirmed that *COI1* expression was significantly reduced in *IRCOI1* plants compared with wt plants (S3B Fig). To determine whether *IRCOI1* plants were impaired in their response to exogenous methyl jasmonate (MeJA), germinated seeds were transferred to agar medium supplemented with 20 or 50 μM MeJA. In the presence of either concentration of MeJA, *IRCOI1* seedlings developed significantly longer roots than wt seedlings. (S3C Fig). Furthermore, MeJA treatment (250 μM) of four-week-old plants induced the expression of the JA-responsive genes *Threonine deaminase* (*TD*) and *Proteinase inhibitor I* (*PI-1*) to higher levels in wt than in *IRCOI1* plants, suggesting that JA responsiveness is altered in *IRCOI1* plants (S3D, E Fig).

Wt, *NahG* and *IRCOI1* transgenic plants were infiltrated with *Agrobacterium* cultures expressing either (-)ssRNA or dsRNA, and inoculated 3 days later with TMV-GFP. At 4 dpi, the size of GFP foci was significantly reduced in dsRNA-expressing patches compared with (-)ssRNA-expressing patches in both *NahG* transgenic (1.9-fold) and WT plants (2.8-fold), whereas no significant difference was observed between dsRNA- and (-)ssRNA-expressing patches in *IRCOI1* transgenic plants (Fig. 6A,B). Consistently, TMV-GFP RNA accumulated less in the patches expressing dsRNA than in (-)ssRNA-expressing patches in wt plants at 5 dpi, whereas viral RNA accumulated at similar levels in patches expressing either (-)ssRNA or dsRNA in *IRCOI1* transgenic plants, as assayed by qRT-PCR (Fig. 6C). Similar levels of both (-)ssRNA and dsRNA were expressed in wt and *IRCOI1* plants (Fig. 6D). The failure of dsRNA to induce a PTI response against TMV-GFP in *IRCOI1* plants was independent of the dsRNA delivery method used. Topical application of *Escherichia coli*-produced dsRNA was unable to reduce the size of TMV-GFP foci compared to the control treatment in *IRCOI1* leaves (S4 Fig).

**Fig 6.**
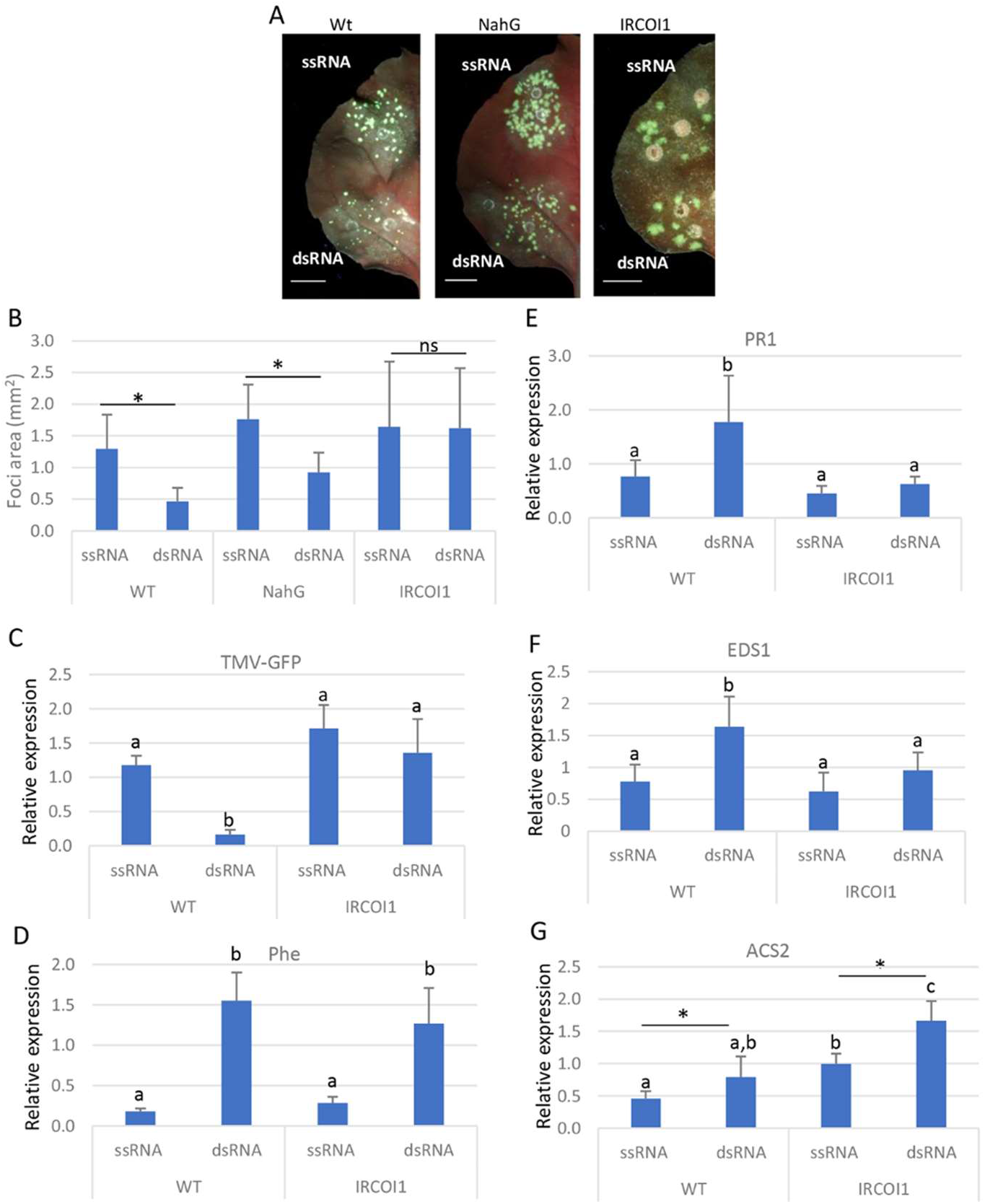
dsRNA-induced PTI is impaired in transgenic plants defective in JA signaling. (A) Leaves of *N. benthamiana* transgenic plants deficient in jasmonic acid perception (*IRCOI1*) or salicylic acid accumulation (*NahG*), as well as wild-type (wt) plants, were infiltrated with *Agrobacterium* cultures containing binary constructs expressing dsRNA derived from PMMoV or (-)ssRNA, as indicated. After 3 days, the infiltrated patches were inoculated with TMV-GFP. The inoculated leaves were examined under UV light at 4 days post-inoculation (dpi). Scale bar denotes 1 cm. (B) Mean numbers ± standard deviations of infection foci on infiltrated patches with the different treatments at 4 dpi. Six patches from 3 plants for each treatment and genotype were analyzed. Asterisks indicate significant differences between treatments (Student’s t-test, *p* < 0.05); ns: not significant. Experiments with *IRCOI1* plants were repeat once more with similar results. (C,D) qRT-PCR was used to analyze the accumulation of TMV-GFP RNA (C) and the phleomycin (Phe) RNA sequence (D) in wt and *IRCOI1* plants at 5 dpi. (E-G) The relative expression levels of the defense-related genes *Pathogenesis-related gene 1* (*PR1*) (E), *Enhanced disease susceptibility 1* (*EDS1*) (F), and the *1-aminocyclopropane-1-carboxylate synthase2* (*ACS2*) (G), were determined by qRT-PCR. Expression of the 18S rRNA gene served as a control. Data represent the means ± standard deviation of 4 replicates, each consisting of a pool of 6 plants that received the same treatment. Different letters indicate significant differences determined by employing Tukey’s test (*p* < 0.05). Experiments with *IRCOI1* plants were repeated once more with similar results.

We then measured by qRT-PCR the expression of the defense-related genes *Pathogenesis-related gene 1* (*PR1*), *Enhanced disease susceptibility 1* (*EDS1*) and *1-aminocyclopropane-1-carboxylate synthase2* (*ACS2*), which has been previously reported to be upregulated by application of dsRNA (Huang et al., 2023; Necira et al., 2024). Figure 6E-G shows a significant induction of *PR1* and *EDS1* in patches expressing dsRNA compared to (-)ssRNA-expressing patches in wt plants. However, both genes accumulated at levels similar to (-)ssRNA-expressing wt patches in *IRCOI1* tissues expressing either (-)ssRNA or dsRNA. The *ACS2* mRNA was upregulated in patches expressing dsRNA compared to (-)ssRNA-expressing patches in both wt and *IRCOI1* plants.

We wonder whether dsRNA-induced callose deposition was altered in *COI1*-silenced plants. After several attempts, *IRCOI1* transgenic plants proved recalcitrant to vacuum infiltration of the dye solution into the leaf tissue. Thus, instead of using *IRCOI1* transgenic plants, we transiently silenced *COI1* by agroinfiltration of a COI1 hairpin construct in wt plants. qRT-PCR analysis showed that *COI1* mRNA levels were significantly reduced by transient expression of *IRCOI1* at 4 days after infiltration (dai) (Fig. 7A). Wt *N. benthamiana* plants were infiltrated with combinations of *Agrobacterium* cultures expressing GUS plus either (-)ssRNA or dsRNA, and *IRCOI1* plus either (-)ssRNA or dsRNA, and inoculated 3 days later with TMV-GFP. At 4 dpi, there was a significant reduction in the size of GFP foci in patches expressing dsRNA compared to (-)ssRNA-expressing patches in control plants, whereas there was no significant difference between patches expressing dsRNA and (-)ssRNA in *COI1-*silenced plants (Fig. 7B,C). In vivo aniline blue staining was performed on patches expressing any of the four combinations mentioned above in the absence of viral infection. Consistent with the lack of a significant reduction in the size of TMV-GFP foci observed in *COI1*-silenced plants, dsRNA treatment did not increase callose deposition at PD in these plants. By contrast, callose accumulation was increased by ca 50% in patches treated with dsRNA compared to (-)ssRNA treatment in control plants (Fig. 7D,E). Interestingly, *COI1* mRNA expression was induced in dsRNA-expressing patches in non-silenced tissue, pointing out that *COI1* is up-regulated by dsRNA treatment (Fig. 7F). The inability of dsRNA to induce a PTI response in *IRCOI1* plants was not limited to TMV-GFP but also extended to TuMV-GFP and TRV-GFP (S5 Fig).

**Fig 7.**
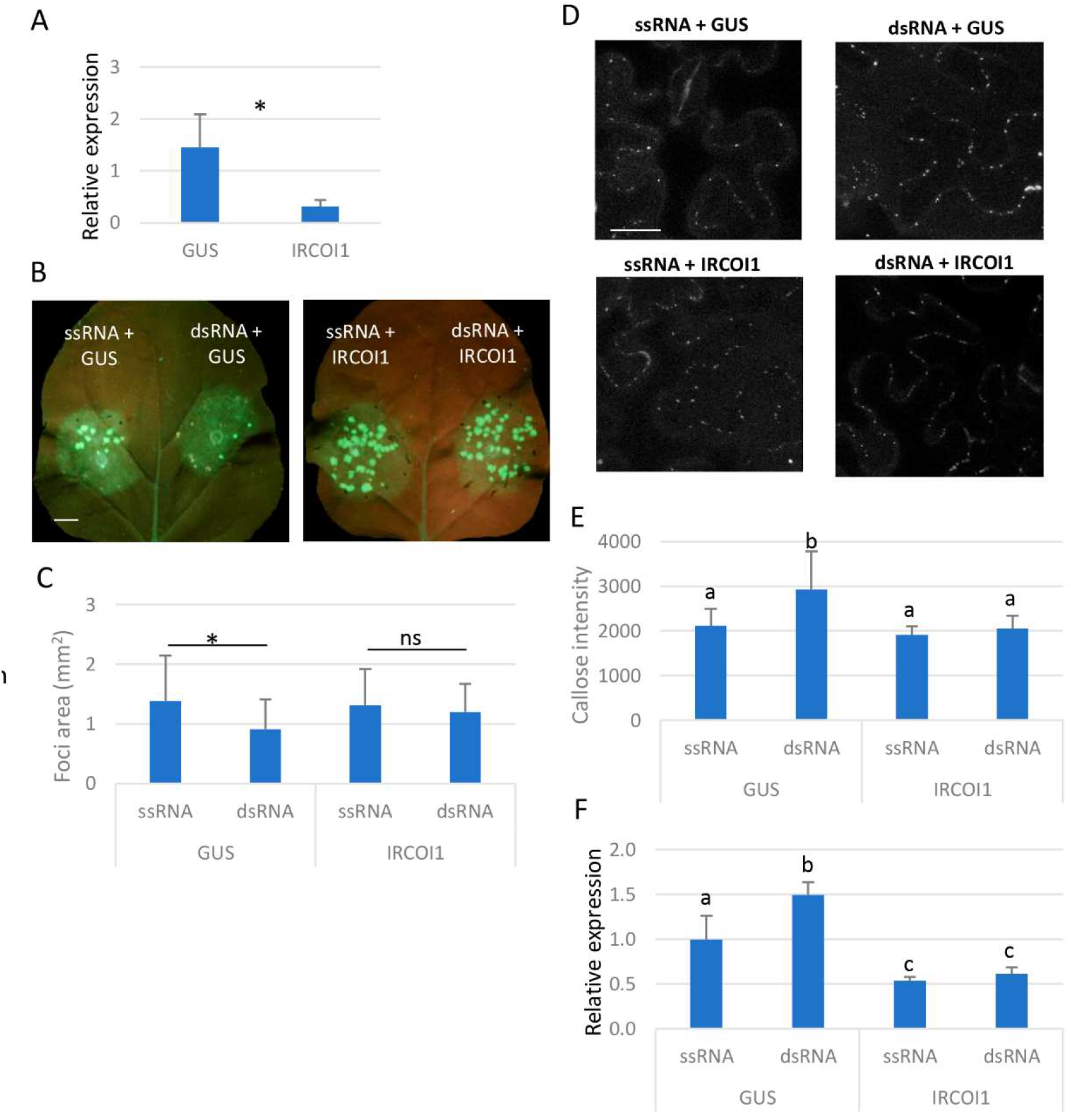
Transient silencing of *COI1* impairs dsRNA-induced restriction of TMV-GFP cell-to-cell movement by preventing callose accumulation. (A) Three-week-old *N. benthamiana* plants were infiltrated with *Agrobacterium* cultures containing binary constructs expressing GUS or hairpin RNA sequences derived from *COI1* (*IRCOI1*). qRT-PCR analysis of expression of *COI1* in silenced plants at 4 days after infiltration (dai). (B) Leaves of *N. benthamiana* were infiltrated with combinations of *Agrobacterium* cultures containing binary constructs expressing dsRNA sequences derived from PMMoV plus either GUS or *IRCOI1*, and expressing (-)ssRNA plus either GUS or *IRCOI1*, as indicated. After 3 days, the infiltrated patches were inoculated with TMV-GFP. The inoculated leaves were examined under UV light at 4 days post-inoculation (dpi). Scale bar denotes 1 cm. (C) Mean numbers ± standard deviations of infection foci on infiltrated patches with the different treatments at 4 dpi. Ten patches from 5 plants for each treatment were analyzed. Asterisks indicate significant differences between treatments (Student’s t-test, *p* < 0.05). ns: not significant. (D) Callose spots at plasmodesmata (PD) were visualized upon aniline blue staining of epidermal cells on patches in response to the different treatments mentioned above, in the absence of viral infection. Scale bar, 10 μm. (E) Relative PD callose content in the different treatments. The mean values of callose intensities in individual PD (>200) measured in three leaf discs taken from three independent biological replicates per each treatment. (F) qRT-PCR analysis of expression of *COI1* mRNA at 4 dai. Different letters indicate significant differences determined by employing Tukey’s test (P < 0.05). Experiments were repeated once more with similar results.

### *COI1* silencing causes transcriptome reprogramming

To investigate whether dsRNA-induced restriction of virus replication is affected in a *COI1*-silenced background, virus-induced gene silencing (VIGS) analysis was conducted in *N. benthamiana* using the TRV system. *N. benthamiana* plants were infiltrated with Agrobacterium cultures expressing either a TRV recombinant vector harboring *COI1* sequences (TRV2:COI1) or the empty vector (TRV2:00) as a control. At 15 dai, *COI1* silencing efficiency was examined in upper, non-infiltrated leaves by qRT-PCR (Fig. 8A). Transcripts of *COI1* accumulated to lower levels in the TRV2:COI1-treated plants compared to the control plants. Then, plants were co-agroinfiltrated in upper leaves with a binary vector expressing TuMV-GFP together with binary vectors expressing either PMMoV (-)ssRNA or dsRNA. Three days after virus inoculation, a reduction in GFP fluorescence intensity was evident in patches expressing dsRNA compared to those expressing (-)ssRNA in both *COI1*-silenced and control plants (Fig. 8B). Consistently, virus accumulation was lower in leaf patches treated with dsRNA than in (-)ssRNA-expressing patches in both types of plants, as assayed by western blot analysis (Fig. 8D). Similar results were obtained using *IRCOI1* transgenic plants (Fig. 8C,D). Thus, *COI1* silencing impairs dsRNA-mediated restriction of viral cell-to-cell movement but it cannot overcome inhibition of viral replication.

**Fig 8.**
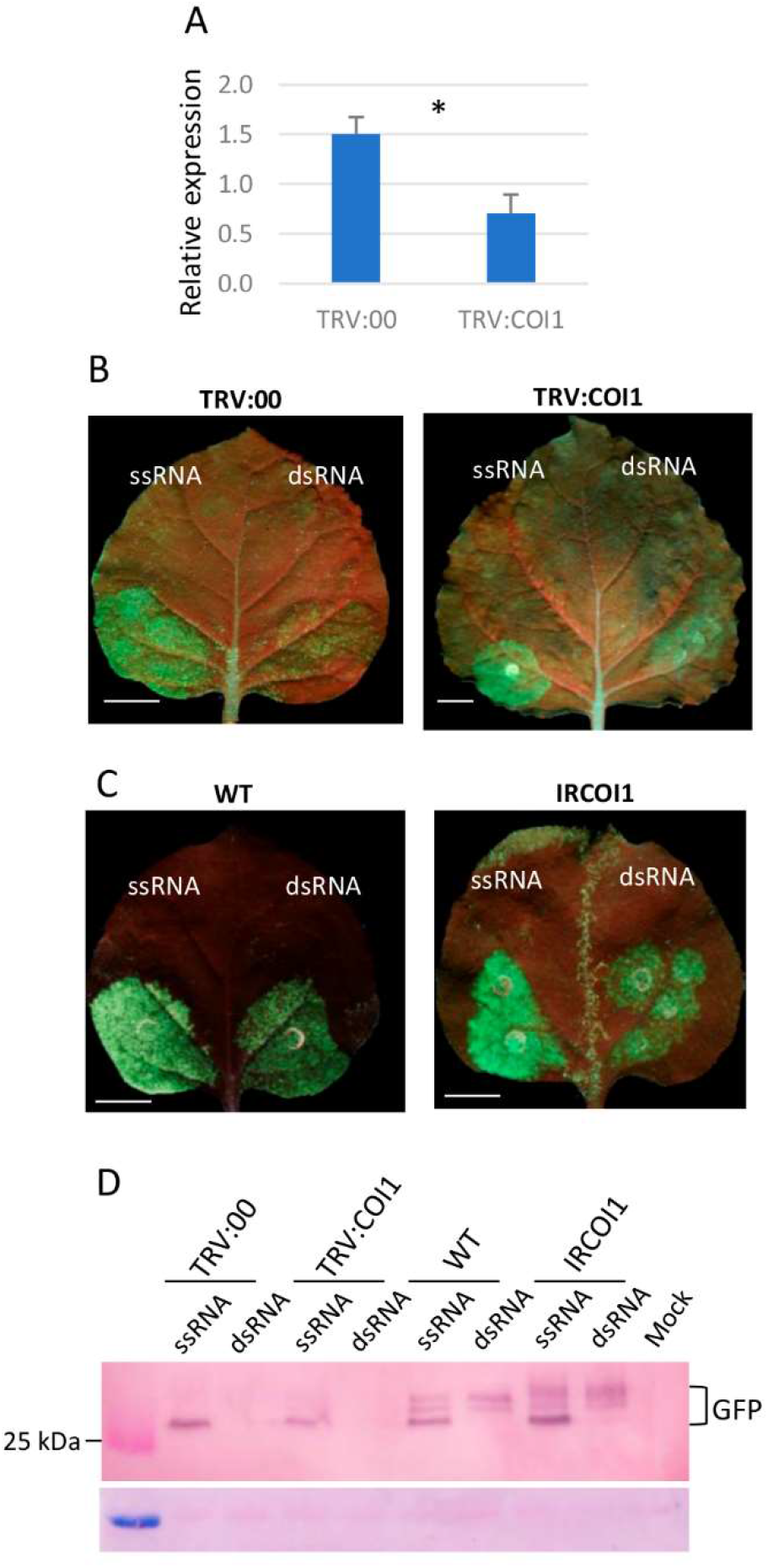
Silencing of *COI1* did not compromise dsRNA-induced restriction of TuMV-GFP replication. (A) Three-week-old *N. benthamiana* plants were infiltrated with *Agrobacterium* cultures containing TRV2:*COI1* or TRV2:00 (vector control). The relative level of expression of *COI1* in silenced plants was estimated by qRT-PCR analysis at 15 days after infiltration (dai). Expression of the 18S rRNA gene served as a control. Data represent the means ± standard errors of three replicates, each consisting of three plants that received the same treatment. Asterisk indicates significant differences between treatments (Student’s t-test, *p* < 0.05). (B) Systemic leaves were infiltrated with *Agrobacterium* cultures containing binary constructs expressing TuMV-GFP together with cultures expressing dsRNA sequences derived from PMMoV or (-)ssRNA at 15 dai, as indicated. The inoculated leaves were examined under UV light at 4 days post-inoculation (dpi). Scale bar denotes 1 cm. (C) Leaves of *N. benthamiana* transgenic plants deficient in jasmonic acid perception (*IRCOI1*) and wild-type (wt) plants were infiltrated with cultures containing binary constructs expressing TuMV-GFP together with cultures expressing dsRNA sequences derived from PMMoV or (-)ssRNA.(D) Western blot analyses of extracts derived from leaf patches in (B) and (C) at 4 dpi, using antibodies against GFP. The upper bands likely correspond to incomplete protein denaturation.

To identify early biological processes involved in COI1-dependent, dsRNA-induced immunity, we conducted a global transcriptome RNA-seq assay with RNA extracted at 4 dai from *N. benthamiana* patches agroinfiltrated with combinations of *Agrobacterium* cultures expressing GUS plus dsRNA (ds_Ctr), and *IRCOI1* plus dsRNA (ds_IRCOI1). Sequencing of 8 transcriptome libraries generated over 748 million mapped reads. On average 93.53% of the clean reads had quality scores at the Q30 level (S1 Table). Principal component analysis (PCA) was used to sort RNA-seq-based transcriptomic data according to gene expression levels. Two principal components explained 70.8% of the overall variance of gene expression profiles (50.6 and 20.2% for principal component 1 (PC1) and principal component 2 (PC2), respectively; Fig. 9A). Data were analyzed using a minimum fold change of 2 (|log_2_ ratio|≥1) and *p*-value ≤ 0.05 as the criteria for differential expression. Comparing ds_IRCOI1 and ds_Ctr datasets, 198 genes were differentially expressed (DEG), among which 147 were expressed at higher levels in ds_IRCOI1 compared to ds_Ctr (S1 Table). The Kyoto Encyclopedia of Genes and Genomes (KEGG) pathway analysis was performed to identify pathways significantly associated with the DEGs (S1 Table). KEGG enrichment analysis (*p* < 0.05) identified two significantly enriched pathways in the ds_IRCOI1 vs ds_Ctr comparison, i.e., Plant–pathogen interaction and Glutathione metabolism (Fig. 9B). The Plant–pathogen interaction pathway comprised seven upregulated DEGs, including those encoding the calcium-binding proteins CML38 and CML46 and the ERF-family transcription factor PTI5. It has been described that CML46, which was upregulated in *COI1*-silenced plants, suppress PTI responses to the pathogen *P. syringae* in Arabidopsis (Lu et al., 2018), whereas PTI5 overexpression enhanced pathogen-induced expression of β-1,3-glucanase in tomato (He et al., 2001). The Glutathione metabolism pathway comprised four DEGs, including those encoding *Glutathione S-transferases* (*GST*s) -of which one was upregulated and two were downregulated- and Ornithine decarboxylase (Table S1). Interestingly, the *AtGST13* mutant has been reported to enhance disease susceptibility toward several fungal pathogens with a concomitant reduction of callose deposition after flg22 treatment (Pislewska-Bednarek et al., 2018).

**Fig. 9.**
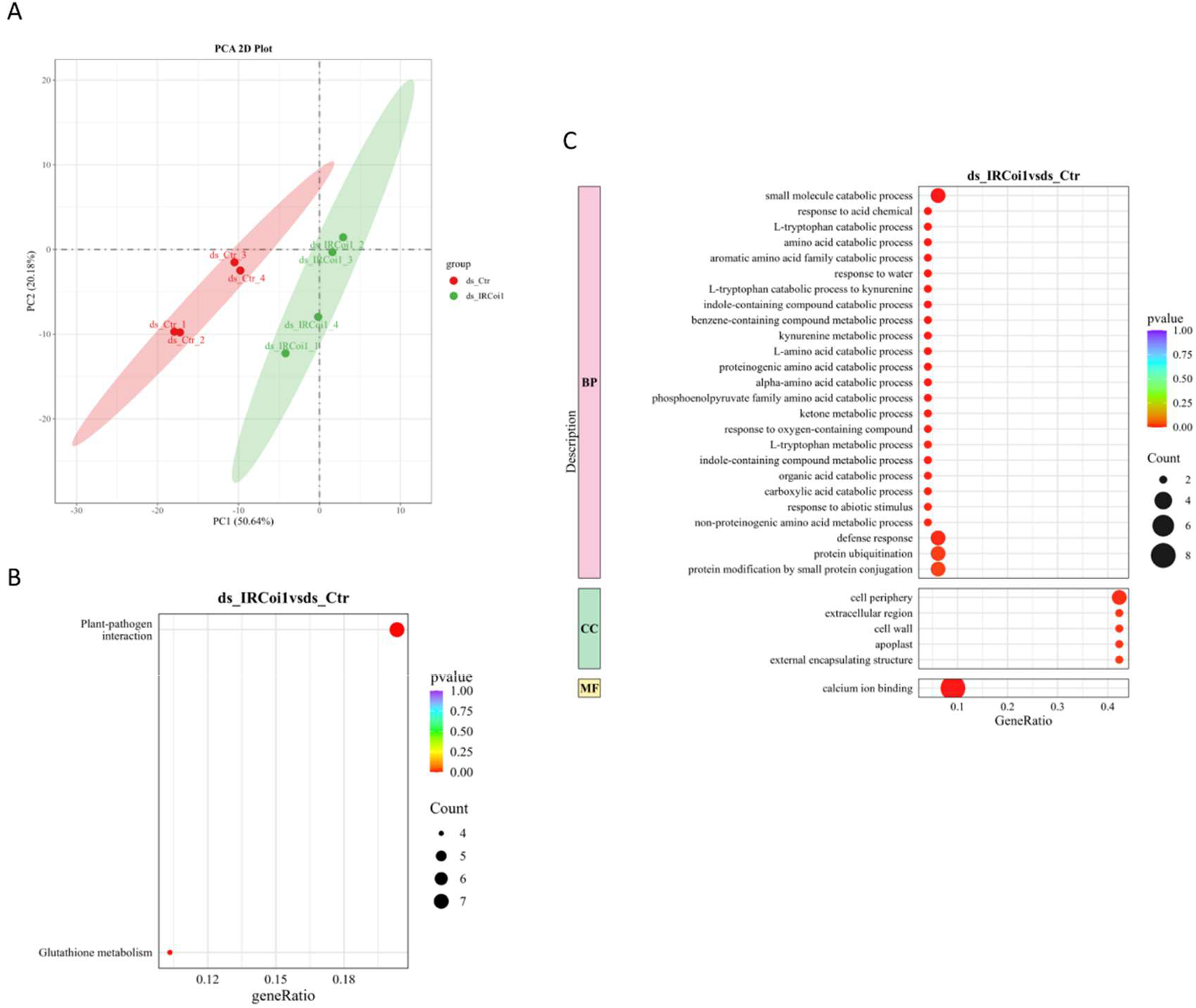
Transcriptional reprogramming associated with dsRNA treatment in a *COI1* silencing background. (A) *N. benthamiana* patches were agroinfiltrated with combinations of *Agrobacterium* cultures expressing GUS plus dsRNA (ds_Ctr) and *IRCOI1* plus dsRNA (ds_IRCOI1). Principal component analysis (PCA) was performed using normalized RNA-seq data at 4 days after infiltration. (B) Kyoto Encyclopedia of Genes and Genomes (KEGG) pathway enrichment analysis (adjusted *p*-value < 0.05) of differentially expressed genes (DEGs) in the ds_IRCOI1 vs ds_Ctr comparison. (C) Gene ontology (GO) analysis of DEGs in the ds_IRCOI1 vs ds_Ctr comparison. Gene ratio is the percentage of total DEGs in the given term. Dot size represents the number of genes annotated to a specific term. Different colors are used to distinguish biological processes, cellular components, and molecular functions.

In addition to KEGG analysis, Gene Ontology (GO) enrichment analysis was performed to identify candidate genes involved in the dsRNA-induced, restriction of viral cell-to-cell movement. Particular attention was given to GO terms associated with Cellular Component and Molecular Function categories (Fig. 9C, S1 Table). The only term overrepresented in the Molecular Function category was Calcium ion binding, which included CML38 and CML46 among others. Regarding the Cellular Component category, several GO terms were overrepresented, with of all DEGs being upregulated. DEGs within Cell periphery and related terms comprise *Xyloglucan endotransglucosylase1* (*XTH1*) and the *Exocyst complex component EXO70H1* involved in autophagy-related, Golgi-independent membrane traffic to the vacuole. *NbXTH*, which was upregulated in *COI1*-silenced plants, has been reported as a positive regulator of cell-to-cell movement of TMV in *N. benthamiana* (Ershova et al., 2025).

## Discussion

In this study, we established an *Agrobacterium*-mediated transient expression assay that enables the induction of dsRNA-mediated PTI in the absence of exogenously applied dsRNA. Transient delivery of a construct directing the production of a hairpin RNA containing nonspecific sequences reduced virus accumulation when co-delivered with binary vectors expressing TuMV-GFP or TRV-GFP, or when followed by mechanical inoculation with TuMV-GFP, TRV-GFP, or TMV-GFP. To date, studies of dsRNA-induced PTI in antiviral immunity have primarily relied on the topical application of either in vivo- or in vitro-synthesized dsRNA (Chi et al., 2026; Huang et al., 2023; Necira et al., 2024; Niehl et al., 2016). As with RNAi triggered by sequence-specific dsRNA, the timing and mode of external application of nonspecific dsRNA may be important factors governing its efficacy as a PTI inducer (Weiss et al., 2025). Thus, simultaneous delivery of binary vectors expressing dsRNA and the virus would increase the likelihood of co-delivery to the same cells. Based on a Poisson distribution model (Carlson et al. 2023), approximately 95% of *N. benthamiana* epidermal cells receiving the viral amplicon are expected also to receive the dsRNA-expressing T-DNA in our experimental conditions. Likewise, mechanical inoculation of viruses onto leaf tissues agroinfiltrated with dsRNA-expressing constructs would favor the presence of both virus and dsRNA within the same cells. Compared with exogenous application of dsRNA, *Agrobacterium*-mediated transient expression provides: i) a sustained intracellular production of dsRNA, rather than a pulse of externally supplied dsRNA, ii) does not depend on uptake/distribution of externally applied dsRNA, and iii) it enables its simultaneous expression with viral replicons or viral proteins. Although *Agrobacterium* itself can activate and modulate plant immune responses through the recognition of bacterial MAMPs and the delivery of virulence-associated effectors (Yang et al., 2023), the comparative study with appropriate controls (ssRNA) allows us to discriminate effects associated with *Agrobacterium*-mediated transient expression from those elicited by nonspecific dsRNA.

It has been reported that the MPs of TMV and other tobamoviruses suppress the dsRNA-induced host response leading to callose deposition at PD (Huang *et al*., 2023). Here, we show that the P25 movement protein of PVX, a virus belonging to a different family (*Alphaflexiviridae*), interferes with nonspecific dsRNA-induced callose deposition, thereby facilitating the movement and accumulation of three distinct viruses, TuMV-GFP, TRV-GFP, and TMV-GFP. Despite their distinct evolutionary origins, PVX P25 and tobamovirus MP proteins share the ability to suppress dsRNA-induced callose deposition at PD, suggesting functional convergence in viral manipulation of this defense barrier (Kumar and Dasgupta, 2021). Previous findings showed that cell-to-cell movement of PVX-GFP was not significantly inhibited by the topical application of nonspecific dsRNA (Necira *et al*., 2024). Thus, the P25 protein encoded by PVX may mask the inhibitory effect of dsRNA-induced PTI on PVX cell-to-cell movement by interfering with callose deposition. These findings support a general model in which viral MPs act as effector proteins that suppress dsRNA-induced PTI by preventing callose accumulation at PD and, consequently, promoting virus movement (Heinlein, 2025). In addition, our approach provides a framework for analyzing both dsRNA-induced responses and their suppression by viral effectors entirely through a transient gene delivery system. The transgenic expression of MP of *Cucumber mosaic virus* has been reported to suppress ROS production and the expression of defense-related genes triggered by bacterial and fungal elicitors of PTI (Kong *et al*., 2018). Together, these findings suggest that viral MPs may broadly function as suppressors of host immune responses by interfering with distinct PTI-associated defense mechanisms.

By using two P25 mutants, T117A and K124E, defective in RNAi suppression, we show that interference with nonspecific dsRNA-induced callose deposition is independent of the RNA silencing suppressor activity of P25. In addition to its role as an RNA silencing suppressor, PVX P25 has been shown to increase the size exclusion limit of PD and to possess RNA helicase activity (Verchot, 2022). Notably, it has been reported that the T117A and K124E mutants differ in their effects on PD gating and helicase activity (Bayne et al., 2005). The T117A mutant is able to alter the PD size exclusion limit, whereas K124E does not. Conversely, K124E retains helicase activity, whereas T117A fails to support this function. Thus, neither PD gating nor helicase activity alone appears to account for the role of P25 in suppressing nonspecific dsRNA-induced callose deposition. Since the P25 T117A mutant did not overcome dsRNA-induced inhibition of viral replication, it is unlikely that P25 interferes with dsRNA perception by a host receptor. Tilsner et al. proposed that P25-mediated remodeling of actin/endomembranes could contribute to PD dilation, facilitating the movement of PVX throughout the PD (Tislner et al., 2012). Previous studies have shown that actin depolymerization affects callose deposition at PD. Actin Depolymerizing Factor 3 (ADF3) together with C-type lectin receptor-like kinase 1 (CLRLK1), act as negative regulators of callose accumulation at PD (Kumar et al., 2024). Callose synthases are delivered to the plasma membrane through vesicular trafficking that depends on the actin cytoskeleton. Therefore, altering actin organization can influence the delivery/localization of callose synthases at PD. Further studies are needed to elucidate how P25 interferes with the antiviral PTI component that restricts virus cell-to-cell movement.

Our experimental approach, based on the transient expression of dsRNA together with viral replicons, enabled the analysis of antiviral defenses elicited by nonspecific dsRNA under conditions in which a substantial proportion of epidermal cells—approximately 70%, according to Carlson et al. (2023)—would receive the viral amplicon, thereby reducing significantly the contribution of cell-to-cell movement to overall virus accumulation. We observed a several-fold reduction in the accumulation of TuMV-GFP and TRV-GFP in patches expressing nonspecific dsRNA compared to those expressing ssRNA. Consistent with the absence of a significant contribution of virus cell-to-cell movement in these assays, co-expression of the P25 T117A mutant did not alleviate the dsRNA-induced inhibition of TuMV-GFP accumulation. Thus, PTI triggered by nonspecific dsRNA restricts virus replication beyond its effects on cell-to-cell movement. Furthermore, we demonstrate that nonspecific dsRNA effectively reduces the accumulation of two viruses in isolated plant cells. Given that key features of PTI signaling are retained in protoplasts (Mueller et al., 2012), this experimental system could provide a useful tool for studying PTI responses elicited by nonspecific dsRNA and identifying potential viral suppressors of PTI-mediated defenses at the cellular level.

A role of JA signaling in the establishment of antiviral PTI induced by nonspecific dsRNA was substantiated from the analysis of plants stably or transiently silenced for *COI1*. *Agrobacterium*-mediated transient expression and topical application of nonspecific dsRNA in *COI1*-silenced tissues had no detrimental effect on virus cell-to-cell movement and/or accumulation of three different viruses. Moreover, unlike in wt plants, transient expression of dsRNA did not induce callose deposition, and the induction of the defense-related genes *PR-1* and *EDS1* was compromised in *COI1*-silenced tissues. COI1 is s a critical component of the JA receptor and a key regulator of JA-responsive gene expression, thereby controlling a broad spectrum of physiological, metabolic, and defense responses (Zhou et al., 2025). Although SA plays a key role in regulating callose deposition at plasmodesmata during pathogen infection (Wang et al., 2013), previous studies have also implicated JA signaling in the regulation of callose accumulation and defense responses. García-Andrade et al. (García-Andrade et al., 2011) showed that potentiation of callose accumulation induced by two necrotrophic fungal pathogens in Arabidopsis required JA perception through COI1. In addition, extracellular self-DNA has been proposed to activate PTI responses through a damage associated molecular pattern-triggered immunity that depends on JA signaling (Zhou et al., 2023). These findings provide support to the involvement of COI1-dependent JA signaling in the regulation of dsRNA-induced callose deposition and the associated restriction of virus cell-to-cell movement.

We aimed to characterize the global transcriptional changes induced by nonspecific dsRNA in a *COI1*-silenced background to identify candidate genes and pathways potentially involved in the dsRNA-mediated restriction of viral cell-to-cell movement. However, the incomplete annotation of the *N. benthamiana* genome and the limited functional characterization of some genes may hinder accurate gene identification, orthology assignment, and functional interpretation. This is particularly relevant for KEGG/GO terms related to complex processes such as PD regulation, callose deposition, and JA signaling (Kurotani et al., 2023). Despite of this, our transcriptomic analysis provided insights into the early molecular responses associated with COI1-dependent, dsRNA-induced antiviral immunity. The differential expression of 198 genes in *COI1*-silenced plants compared with control plants suggests that JA signaling contributes to shaping the transcriptional response to nonspecific dsRNA.

Among the DEGs upregulated in *COI1*-silenced plants, CML38 and CML46 are of particular interest because calcium signaling and calcium-binding proteins have been involved in the dsRNA-induced callose deposition at PD (Huang et al, 2023). Calcium-binding proteins, including calmodulins and CMLs, play complex and sometimes opposing roles in PTI, acting as calcium sensors that decode pathogen-induced changes in cytosolic calcium concentrations and translate them into downstream defense responses (Zeng et al., 2023). Depending on the specific protein, its targets, and the context of activation, these sensors can positively regulate immunity by modulating defense-related gene expression, protein kinase activity, and other signaling processes, or negatively regulate PTI by suppressing immune signaling and limiting defense activation. For example, CML46 and CML47 have been shown to negatively regulate PTI responses against *P. syringae* in Arabidopsis, demonstrating that calcium-binding proteins do not invariably act as positive regulators of defense (Lu et al., 2018). Notably, CML41, a Calmodulin-like protein positively regulating dsRNA-induced callose accumulation at PD (Huang et al, 2023), was downregulated in *COI1*-silenced plants (log_2_(Fold-change) = -0.51, *P* = 0.015), although this change did not meet the differential expression threshold applied in S1 Table. Similarly, the upregulation of *PTI5*, an ERF-family transcription factor that stimulates pathogen-induced *β-1,3-glucanase* expression in tomato (He et al., 2001), suggests an enhanced degradation of callose in *COI1*-silenced plants, although its contribution to dsRNA-induced immunity remains to be established.

The enrichment of glutathione metabolism among the DEGs present in the ds_IRCOI1 vs ds_Ctr comparison suggested that cellular redox homeostasis may contribute to COI1-dependent, dsRNA-induced antiviral immunity. Changes in redox status and ROS levels have been linked to changes in PD permeability and callose accumulation (Benitez-Alfonso et al., 2011; Vu et al., 2023). Consistent with this view, *GSTU13* has been reported to positively contribute to pathogen resistance and flg22-induced callose deposition in Arabidopsis (Pislewska-Bednarek et al., 2018). Although glutathione S-transferases may influence these processes indirectly through their roles in redox regulation and stress responses, a direct involvement of the identified GSTs in PD function or callose turnover mediated by nonspecific dsRNA remains to be established.

GO enrichment analysis also highlighted several Cellular component terms associated with cell periphery, suggesting that cell periphery-associated processes may contribute to the altered response of *COI1*-silenced plants to nonspecific dsRNA. Notably, the upregulation of *XTH1* is intriguing because *NbXTH* has been implicated in promoting TMV cell-to-cell movement (Ershova et al., 2025). It has been argued that NbXTH accumulation results in cell wall loosening, which in turn creates favorable conditions for TMV local spread. Overall, these findings identify candidate genes and processes associated with the COI1-dependent response to nonspecific dsRNA, providing a basis for further investigation of the molecular links between JA signaling, calcium-dependent regulation, redox homeostasis, and the restriction of viral cell-to-cell movement.

## Materials and Methods

### Binary vectors

Binary vectors encoding for TMV-GFP (Lindbo, 2007), TuMV-GFP (Dunoyer et al, 2004), and TRV-GFP (Donaire et al., 2008) have been described previously. The cDNA construct expressing a hairpin RNA containing PMMoV polymerase gene sequences spanning from nucleotides 3411 to 4388 in the binary vector pBI121 has been previously described (Tenllado et al., 2003). For comparative purpose, two additional constructs containing linear PMMoV sequences in the sense ([+]ssRNA) or antisense ([-]ssRNA) orientations cloned in pBI121 were used. All the three constructs contain a bacterial gene conferring resistance to phleomycin (Phe) designed to improve the stability of the hairpin construct.

TRV polymerase gene sequences spanning from nucleotides 3906 to 5400 were amplified with gene-specific primers (S1 Table) from TRV RNA1 and cloned in the sense ([+]ssRNA) or antisense ([-]ssRNA) orientations in pCAMBIA2300 under the control of the *Cauliflower mosaic virus* 35S promoter. The plasmids obtained were introduced separately into *A. tumefaciens* strain LBA 4404.

T7-tagged versions of wild-type P25 or P25 mutants T117A and K124E were constructed via PCR as described (Aguilar et al., 2015), using primers that introduced the T7 epitope at the C-terminus of the P25 coding sequence, and transferred to pCAMBIA2300.

### Plant material and agroinfiltration

*N. benthamiana* plants were grown in a growth chamber with a 16 h-light/8h-dark cycle at 25°C. The transgenic *N. benthamiana* plants expressing the salicylate hydroxylase gene (*NahG*) have been described previously (Ying et al., 2010). *COI1-*silenced transgenic *N. benthamiana* plants (*IRCOI1*) were obtained as described (García-Marcos et al., 2013). *A. tumefaciens* cultures were grown to exponential phase in Luria–Bertani (LB) medium with antibiotics at 28°C. For infiltration of RNA-expressing constructs and viral proteins, bacterial cultures were diluted to a final optical density of 0.5 at 600 nm and equal volumes of each suspension were combined. For infiltration with viral amplicons, bacterial cultures were diluted to a final optical density of 0.1. Infiltration of the mixtures was performed into leaves of *N. benthamiana* plants as described (Tenllado and Díaz-Ruíz, 2001).

According to the Carlson et al. (2023) Poisson model, the probability that a cell receives at least one T-DNA is:

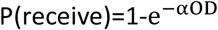

Where the constant term *α* is defined as the mean number of unique T-DNA products delivered by an *Agrobacterium* strain to a given plant cell per infiltration OD_600_ for a given pavement cell, and assuming independent delivery of the *Agrobacterium* strains: dsRNA culture: OD_600_ = 0.25; virus culture: OD_600_ = 0.10; P25/GUS culture: OD_600_ = 0.25. Total OD_600_ = 0.60.

The percentage of virus-receiving cells that also receive dsRNA is determined by the dsRNA delivery probability:

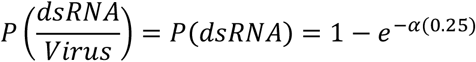

The virus OD_600_ does not enter this conditional probability, provided delivery events are independent. Carlson et al. use α = 12 as the representative value for OD_600_ = 0.2/strain, total OD_600_ = 0.6, which is a good proxy for our experimental conditions.

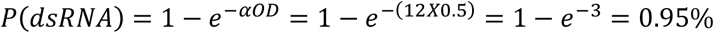

Regarding the percentage of cells receiving the virus, using the same Poisson model with α = 12 and the virus culture at OD_600_ = 0.10:

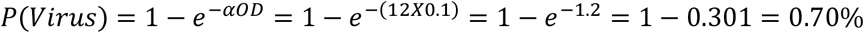

So, **≈**70% of cells are expected to receive the viral amplicon.

For mechanical inoculation, plants were inoculated by rubbing agroinfiltrated patches with plant sap infected with TMV-GFP (1:10 dilution in sodium phosphate buffer 0.02 M, pH 8), TuMV-GFP (1:5), or TRV-GFP (1:5), as described previously (Necira et al., 2024). Viral movement within the inoculated leaves was quantified by measuring the size of GFP-expressing infection foci using ImageJ (https://imagej.net/ij).

In silencing suppression assays, a free GFP reporter gene expressed from pCAMBIA2300 was co-infiltrated with either the pCAMBIA1305.1 vector (GUS) or with pCAMBIA2300 expressing T7-tagged versions of wt P25 or P25 mutants. When possible, all of the treatment types to be compared were infiltrated on a single leaf. To minimize the effects of interleaf variability, all of the leaf disks from different plants corresponding to the same treatment type were pooled.

### Callose staining

Callose deposition was assessed as described previously (Huang *et al*., 2022). Briefly, individual leaf discs from patches infiltrated 2 days earlier with different *Agrobacterium* combinations were vacuum-infiltrated with 0.1% aniline blue prepared in 50 mM potassium phosphate buffer (pH 8.0). As a control treatment, GUS-infiltrated patches were soaked with 0.1% aniline blue supplemented with 1 µM flg22 (MedChemExpress, NJ, USA). Fluorescence was imaged 30 min after staining using a Leica TCS SP8 STED 3X confocal microscope (Wetzlar, Germany) equipped with Leica Application Suite X software. Samples were excited with a 405-nm diode laser, and emission was collected at 430–490 nm using a HC PL APO 40×/1.30 Oil CS2 objective. Images were acquired at 8-bit resolution. Callose fluorescence intensity was quantified using ImageJ and the calloseQuant plugin (Huang *et al*., 2022). Callose deposits were quantified from 5–6 (Fig1) or 7-9 (Fig 4) images obtained from three leaf discs per plant, using at least three plants per treatment.

### dsRNA production

pL4440-CP contains a 902-bp sequence of the PVY coat protein gene flanked by T7 promoters on both sides of the cloning site (Necira et al., 2024). For the in vitro synthesis, pL4440-CP was linearized in independent reactions with *BamH*I and *Hind*III. Once linearized, the digested plasmids were purified and used as templates for transcription of (+) and (-)ssRNAs using the T7 Transcription kit (Roche Molecular Biochemicals, Barcelona). For DNA removal, the synthetic ssRNAs were treated with DNAse I (Sigma-Aldrich, MO, USA) for 10 min at 37 °C and recovered by precipitation. The ssRNA preparations were mixed and heated at 95°C for 2 min, followed by annealing at room temperature for 30–60 min. DsRNA quantitation was done with the ND-1000 spectrophotometer (Nanodrop, Wilmington, DE, USA) and examined in 1% agarose gels. Five µg of dsRNA or (-)ssRNAs in 10 ul combined with a 10 µl-dose of TMV-GFP infected sap was applied to two leaves previously dusted with carborundum as abrasive (Carlo Erba, Barcelona, Spain).

For in vivo dsRNA production, *E. coli* HT115(DE3) carrying L4440-CP was grown in LB medium supplemented with ampicillin (500 µg/ml) and tetracycline (12.5 µg/ml) for 16 h at 37°C. Cultures were then diluted 75-fold in the same medium and grown to an OD₆₀₀ of 0.5. T7 RNA polymerase was induced with 10 µM IPTG, followed by incubation for 2 h at 37°C with shaking. Cells were harvested by centrifugation (4,000 × g, 15 min) and resuspended in ammonium acetate 1 M (Tenllado et al., 2003 bis). Total nucleic acids were extracted by phenol–chloroform treatment followed by ethanol precipitation and adjusted to 0.5 µg/µl with distilled water. Nucleic acids from non-induced HT115(DE3)/L4440-CP cultures were used as a negative control. A 10 µl-dose of TMV-GFP infected sap combined with an equal volume of dsRNA or non-induced *E. coli* extract was applied to two leaves previously dusted with carborundum.

### Protoplast isolation and virus inoculation

*N. benthamiana* protoplasts were prepared from 5–6-week-old plants. Leaves were surface-sterilized by immersion in 10% bleach for 15 min, followed by three washes with sterile water. Leaf strips were digested with 0.5% Cellulase Onozuka R10 and 0.15% Macerozyme R10 (Duchefa, Haarlem, The Netherlands) in 12% mannitol containing 6 mM CaCl₂ (pH 5.7) for 18 h at 25°C in the dark, as described previously (Ammirato et al., 1984). Protoplasts were counted using a hemocytometer and adjusted to ≈85000 cells/mL in ice-cold transfection buffer (12% mannitol, 6 mM CaCl₂, 80 mM KCl, pH 7.2). Aliquots (100 µL) were gently mixed with 5 µg of binary vector expressing TuMV-GFP or TRV RNA1 and 5 µg of PVY-derived dsRNA or (-)ssRNA. For mock transfections, nucleic acids were replaced with equivalent volumes of sterile water. Transfection was initiated by adding 110 µL of PEG–calcium solution (40% PEG 4000, 0.2 M mannitol, 100 mM CaCl₂). After gentle mixing by tapping, samples were incubated for 10–15 min at room temperature. Transfection was terminated by adding 400 µL of 12% mannitol/6 mM CaCl₂. Protoplasts were recovered by centrifugation (100 × g, 2 min), resuspended in 1 mL of 12% mannitol/6 mM CaCl₂, and incubated for 20 h before further analysis.

### qRT-PCR and protein gel blot analysis

Total RNA was extracted from infiltrated leaves at 2 and 5 dpi as described (Pacheco et al., 2012). qRT-PCR for virus accumulation and gene expression was performed with gene-specific primers (S1 Table) as described by García-Marcos and associates (). All reactions were done in triplicate with two replicates of each sample in each run. At least three independent biological replicates for each treatment were used for qRT-PCR analyses and the values obtained averaged. The amplification efficiency was determined for each primer pair with a 3-fold serial dilution of an RNA sample (∼100 ng RNA/μl) with at least five dilutions points. The relative quantification of PCR products was calculated by the comparative ΔΔCt (cycle threshold) method. 18S rRNA was chosen for normalization because of its similar level of expression across all treatments.

Proteins were analyzed by western blot as described in Makki et al. (2026). T7-tagged P25 was detected with horseradish peroxidase conjugated anti-T7 antibody (Novagen) (1:5000 dilution). The GFP or actin protein bands were detected using a rabbit polyclonal antibody to GFP (1:5000 dilution), kindly provided by Professor Lesley Torrance (The James Hutton Institute, UK), or antiserum to actin (1:3000 dilution) (AS13 2640 Agrisera), respectively. Detection was performed using either the ECL system (Amersham Biosciences, Little Chalfont, UK) or with SigmaFast^TM^ BCIP/NBT substrate tablets (Sigma-Aldrich). The detected protein bands were subjected to densitometry analysis using ImageJ software.

### VIGS assays and transient silencing

A 174-bp fragment (nucleotides 1292 to 1465) of *N. benthamiana COI1* was amplified and introduced into the binary vector pTRV2 to generate pTRV2:*COI1*, as described (García-Marcos, 2013). The pTRV1 vector and the pTRV2 vector and its derivative were separately transformed into *A. tumefaciens* strain GV2260. *N. benthamiana* leaves were infiltrated with *A. tumefaciens* cultures as described (Tenllado and Díaz-Ruíz, 2001). Total RNA was extracted from upper, non-inoculated leaf at 15 dai. Silencing of *COI1* mRNA was detected by qRT-PCR as described above. Primers that anneal outside the region targeted for silencing were used to ensure that the endogenous gene is tested.

Transient silencing of *COI1* in *N. benthamiana* was carried out by transferring the 174-bp fragment from pTRV2:*COI1* into the binary destination vector pH7GWIWG2(II) (Karimi et al., 2002), as described (García-Marcos, 2013). The *IRCOI1* construct carries a cassette containing a 35S promoter-driven *COI1* sequence in both sense and antisense orientations, separated by an intron. RNA splicing of a transcript derived from this construct would generate a dsRNA that functions as a potent silencing inducer (Smith et al., 2001).

### Hormone treatment

Four-week-old wt and *IRCOI1* transgenic plants were sprayed with MeJA (250 μM) (Sigma-Aldrich) or water for three consecutive days. After spraying, samples were harvested at 24 h.

To determine the MeJA sensitivity of seedlings, wt and *IRCOI1* seedlings were grown on MeJA-containing agar. As MeJA inhibits germination of the seeds, they were pre-germinated in agar plates containing Murashige and Skoog (MS) medium. After 7 days, the seedlings were transferred to freshly prepared MS agar plates containing 20 and 50 μM MeJA or no hormone. Root length was measured after 15 days using ImageJ software.

### Library Preparation for Transcriptome Sequencing

Four independent biological replicates were used to monitor differences in gene expression between treatments, each replicate consisting of a pool of 12 treated leaves. The integrity and quality of the total RNA were checked using NanoDrop 2000 Spectrophotometer (Thermo Scientific, Waltham, MA, USA). The transcriptome libraries, sequencing and bioinformatics analysis were performed at Novogene in Germany; mRNA was purified from total RNA using poly-T oligo-attached magnetic beads. After fragmentation, the first strand cDNA was synthesized using random hexamer primers, followed by the second strand cDNA synthesis. The library was checked with Qubit and real-time PCR (Rotor-Gene Q thermal cycler, Qiagen, Hilden, Germany) for quantification and bioanalyzer for size distribution detection. Quantified libraries were pooled and sequenced on Illumina NovaSeq PE150 Platform according to effective library concentration and data amount. The clustering of the index-coded samples was performed according to the manufacturer’s instructions. After cluster generation, the library preparations were sequenced and paired-end reads were generated. Raw reads were firstly processed through in-house Perl scripts. In this step, clean reads were obtained by removing reads containing adapter, reads containing ploy-N and low-quality reads from raw data. At the same time, Q30 of the clean data were calculated (Table S1). All the downstream analyses were based on the clean data with high quality.

The *N. benthamiana* reference genome and gene model annotation files were downloaded from the SGN ftp site (https://nbenthamiana.jp/downloads) directly. An index of the reference genome was built using Hisat2 v2.2.1 and paired-end clean reads were aligned to the reference genome using Hisat2 v2.0.5. FeatureCounts v2.1.1 was used to count the reads numbers mapped to each gene. The expected number of Fragments Per Kilobase of transcript sequence per Millions of base-pairs sequenced (FPKM) of each gene was calculated based on the length of the gene and reads count mapped to this gene. RNA-seq data were uploaded to the Gene Expression Omnibus (GEO, http://www.ncbi.nlm.nih.gov/geo). The accession number is GSE352041.

### Enrichment Analysis of Differentially Expressed Genes

Differential expression analysis of the assayed treatments (four replicates per treatment) was performed using the DESeq2 R package (v1.42.0) (Anders and Huber, 2010). The resulting *p*-values were adjusted using the Benjamini and Hochberg’s correction for controlling the false discovery rate. Genes with a |log_2_(Fold-change)| ≥ 1 and adjusted *p-*value ≤ 0.05 were assigned as differentially expressed. Fold-change calculations were performed for paired-comparisons made between treatments.

KEGG and GO enrichment analysis of DEGs was implemented by the clusterProfiler R package (v4.8.1), in which gene length bias was corrected. Terms and pathways with adjusted *p* value less than 0.05 were considered significantly enriched.

### Statistical analysis

All representative images reflect a minimum of three biological replicates. The exact number of biological replicates performed per experiment are indicated in the figure legends. All statistical analyses were performed using the statistical software IBM SPSS Statistics v.25 (IBM Corp., Armonk, NY). For each experiment, samples were assessed for normality via the Shapiro– Wilk test and for equality of variances using Levene’s test. For comparisons between pairs of means (pairwise comparisons), Student’s *t*-test was employed. Details about the statistical approach used can be found in the figure legends.

## Acknowledgments

The research was funded by grants PID2022-137691OB-I00 and PID2025-172502OB-I00 from the Spanish Ministry of Science and Innovation (MCIN/AEI/10.13039/501100011033), and by “ERDF A way of making Europe”. The funder did not play any role in the study design, data collection and analysis, decision to publish, or preparation of the manuscript.

## Author contributions

**Conceptualization:** Francisco Tenllado.

**Formal analysis:** Khouloud Necira, Ifigeneia Triantafyllou, Celia González.

**Funding acquisition:** Francisco Tenllado, Tomas Canto.

**Investigation:** Khouloud Necira, Ifigeneia Triantafyllou, Celia González, Francisco Tenllado.

**Methodology:** Khouloud Necira, Francisco Tenllado.

**Project administration:** Francisco Tenllado, Tomas Canto.

**Software:** Khouloud Necira, Ifigeneia Triantafyllou, Celia González.

**Supervision:** Khouloud Necira, Francisco Tenllado.

**Writing – original draft:** Francisco Tenllado.

**Writing – review & editing:** Khouloud Necira, Tomas Canto, Francisco Tenllado.

