## Supplemental Figures for "Pattern-triggered immunity induced by dsRNA against plant viruses involves two distinct branches, one of which depends on *Coronatine insensitive1* (*COI1*)"

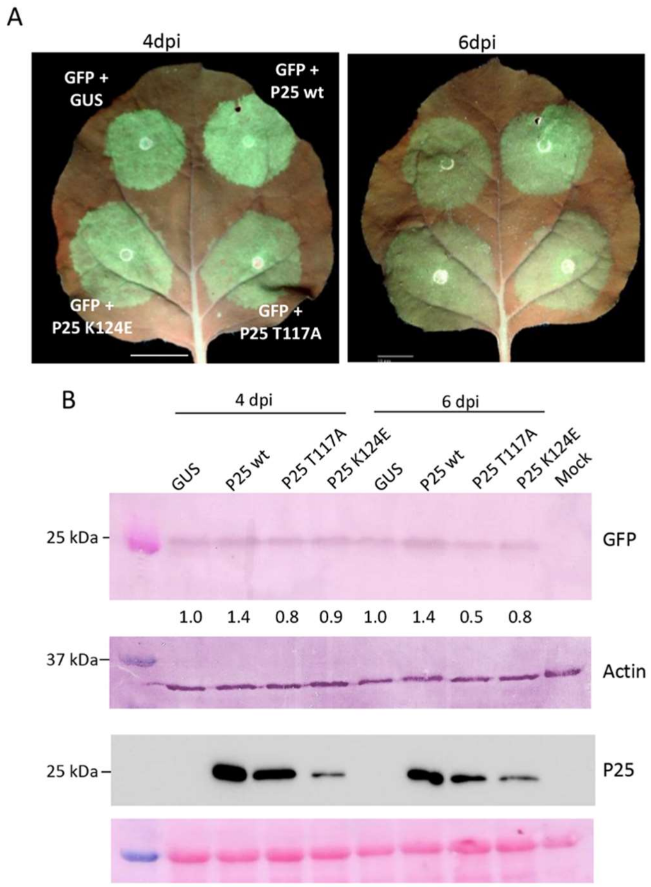


**S1 Fig.** PVX P25 mutants defective in RNA silencing. (A) Leaves of *N. benthamiana* were infiltrated with combinations of *Agrobacterium* cultures containing binary constructs expressing GFP plus either GUS, T7-tagged versions of PVX P25 wt, or P25 mutants T117A and K124E, as indicated. Leaves were photographed at 4 and 6 days post-agroinfiltration (dpa). Scale bar denotes 1 cm. (B) Western blot analyses of extracts derived from leaf patches at 4 and 6 dpa, using antibodies against GFP (upper panel), actin (middle panel) as a loading control, or the T7 epitope (lower panel). The intensity of the GFP bands, normalized for the loading controls, was quantified by densitometry analysis. The value for the band in leaf patches infiltrated with GFP plus GUS was set at 1 and other data calculated relative to this value. The panel below the western blot shows the Ponceau S-stained membrane after blotting. Experiments were repeated once more with similar results.

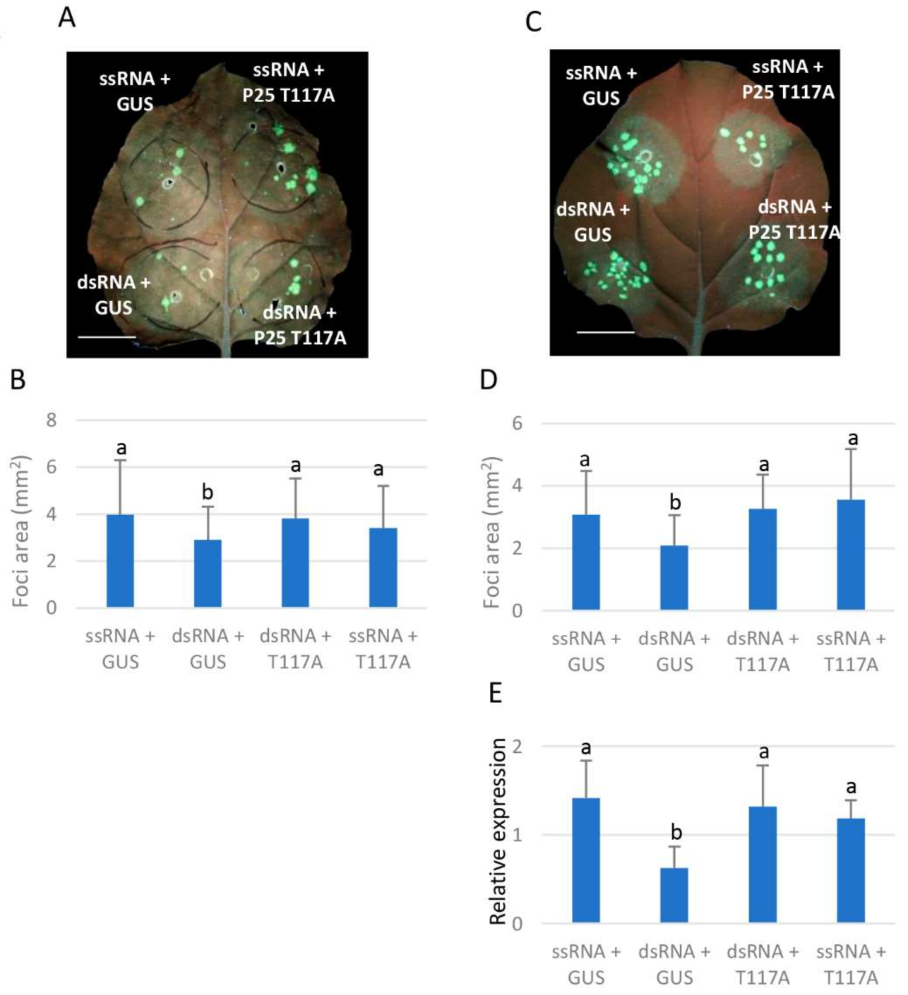


**S2 Fig.** PVX P25 compromises dsRNA-induced restriction of TuMV-GFP and TRV-GFP cell-to-cell movement. (A,C) Leaves of *N. benthamiana* were infiltrated with combinations of *Agrobacterium* cultures containing binary constructs expressing dsRNA sequences derived from PMMoV plus either GUS or T7-tagged P25 T117A, and expressing (-)ssRNA plus either GUS or T7-tagged P25 T117A, as indicated. After 3 days, the infiltrated patches were inoculated with TuMV-GFP (A) or TRV-GFP (C). The inoculated leaves were examined under UV light at 4 (A) or 2 (C) days post-inoculation (dpi). Scale bar denotes 1 cm. (B,D) Mean numbers ± standard deviations of infection foci from TuMV-GFP (B) or TRV-GFP (D) on infiltrated patches with the different treatments at 4 and 2 dpi, respectively. Eight patches from 4 plants for each treatment were analyzed. Different letters indicate significant differences determined by employing Dunnett's T3 test ($P < 0.05$). (E) qRT-PCR was used to analyze the accumulation of TRV-GFP at 4 dpi. Expression of the 18S rRNA gene served as a control. Data represent the means ± standard deviation of at least 3 replicates, each consisting of a pool of 6 plants that received the same treatment. Different letters indicate significant differences determined by employing Duncan's test ($P < 0.05$).

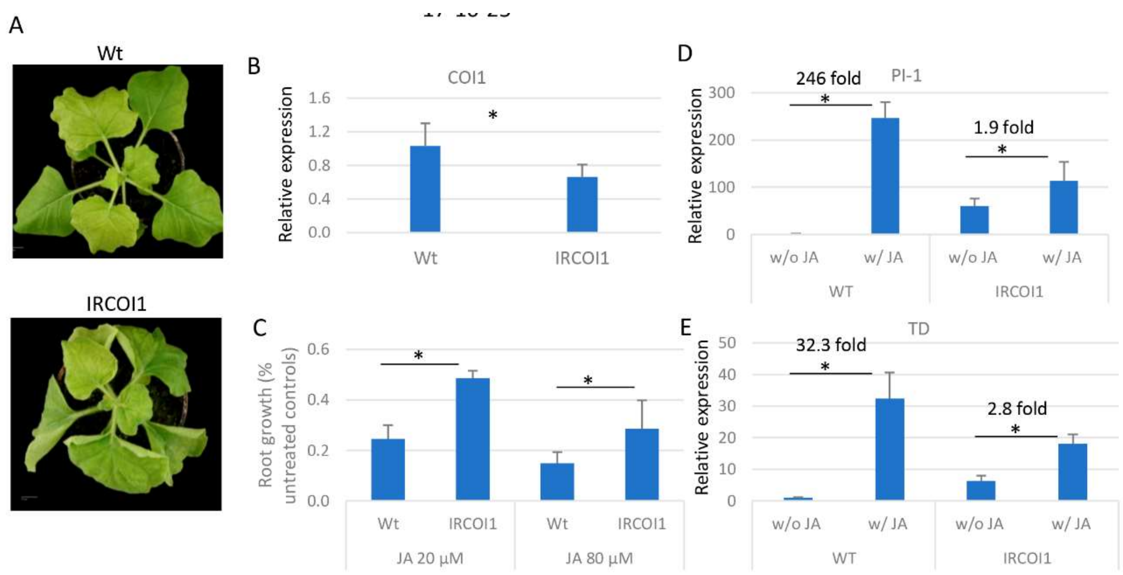


**S3 Fig**. *IRCOI1* transgenic plants are partially insensitive to JA signaling. (A) Four-week-old *IRCOI1* and wt plants . (B) Silencing of *COI1* transcripts in *IRCOI1* as measured by qRT-PCR. (C) Wild-type (wt) and *IRCOI1* seedlings were grown on MS medium containing MeJA at different concentrations (20 and 80 µM). Root growth under MeJA treatment is presented relative to growth on MeJA-free control medium (%). (D) MeJA (250 µM) or mock solution as a control was applied to wt and *IRCOI1* plants, and the expression of *Proteinase inhibitor I* (*PI-1*) and *Threonine deaminase* (*TD*) mRNAs were measured by qRT-PCR. Expression of the 18S rRNA gene served as a control. Data represent the means ± standard deviation of 4 replicates, each consisting of a pool of 6 plants that received the same treatment. Asterisks indicate significant differences between treatments (Student's t-test, $p < 0.05$).

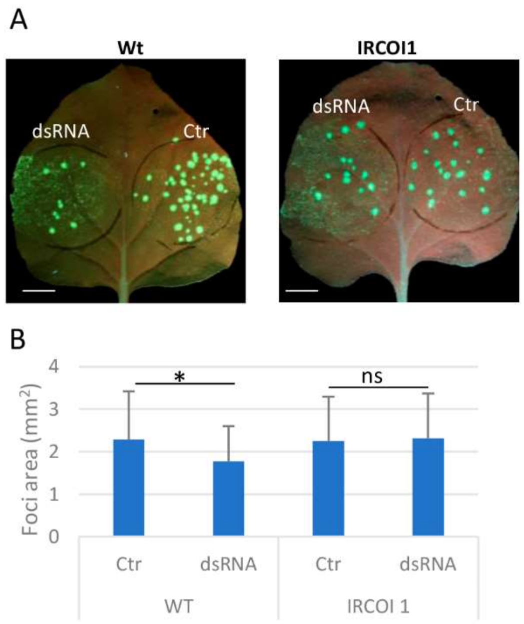


**S4 Fig**. Topical application of *Escherichia coli*-produced dsRNA did not restrict TMV-GFP cell-to-cell movement in *IRCOI1* plants. (A) Leaves of wild-type (wt) and transgenic COI1 (*IRCOI1*) *N. benthamiana* plants were inoculated with mixtures of TMV-GFP combined with either nucleic acid extracts prepared from *E. coli* accumulating dsRNA from PVY (dsRNA) or non-induced control extract (Ctr). Scale bar denotes 1 cm. (B) Mean numbers ± standard deviations of infection foci on patches with the different treatments at 4 dpi. Ten patches from 5 plants for each treatment and genotype were analyzed. Asterisks indicate significant differences between treatments (Student's t-test, $p < 0.05$); ns: not significant.

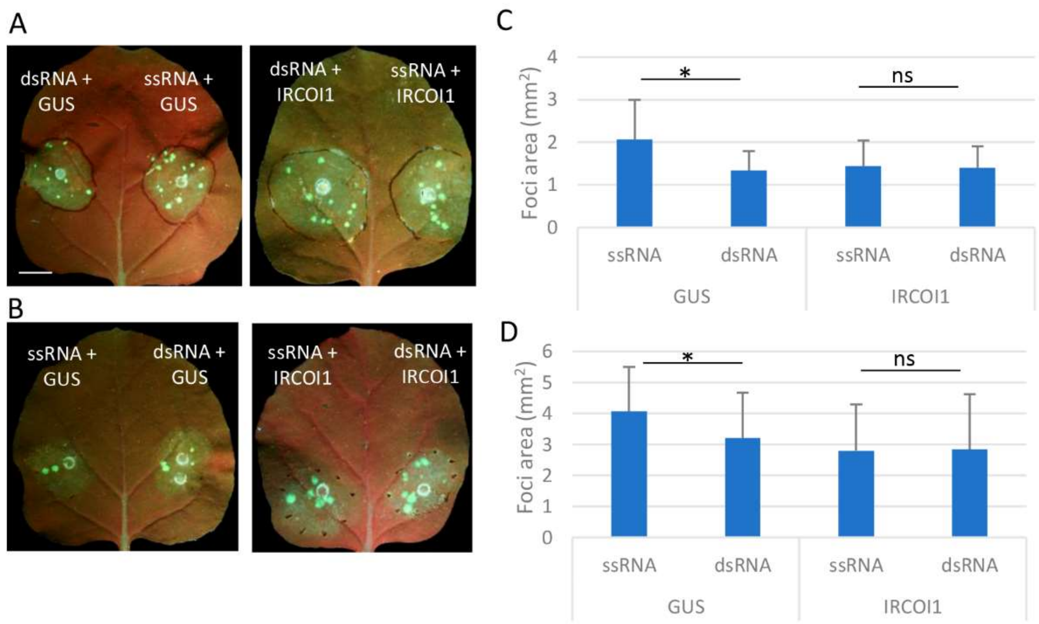


**S5 Fig**. Transient silencing of *COI1* also compromises dsRNA-induced restriction of TuMV-GFP and TRV-GFP cell-to-cell movement. (A,B) Leaves of *N. benthamiana* were infiltrated with combinations of *Agrobacterium* cultures containing binary constructs expressing dsRNA sequences derived from PMMoV plus either GUS or hairpin RNA sequences derived from *COI1* (*IRCOI1*), and expressing (-)ssRNA plus either GUS or *IRCOI1*, as indicated. After 3 days, the infiltrated patches were inoculated with TuMV-GFP (A) or TRV-GFP (B). The inoculated leaves were examined under UV light at 4 (A) or 2 (B) days post-inoculation (dpi). Scale bar denotes 1 cm. (C,D) Mean numbers ± standard deviations of infection foci from TuMV-GFP (C) or TRV-GFP (D) on infiltrated patches with the different treatments at 4 and 2 dpi, respectively. Six patches from 3 plants for each treatment were analyzed. Asterisks indicate significant differences between treatments (Student's t-test, $p < 0.05$). ns: not significant.
